# The population structure of invasive *Lantana camara* is shaped by its mating system

**DOI:** 10.1101/2024.10.22.619585

**Authors:** P. Praveen, Rajesh Gopal, Uma Ramakrishnan

## Abstract

Over the last century, invasive species have emerged as an important driver of global biodiversity loss. Many invasive species have low genetic diversity in the invaded habitats, owing to the demographic bottleneck during introduction. *Lantana camara* is one of the hundred most problematic invasive species globally. Despite its ecological importance in many countries, our understanding of the genetic diversity patterns of this plant remains poor. Previous studies hypothesize that invasive *L. camara* is a species complex with a hybrid origin, though this has never been tested. We investigated the population genetic patterns of *L. camara* by sampling 359 plants that represented a spectrum of flower colour variants across 36 locations, spanning most of the biogeographic regions across India. Analyses of the population structure using 19,008 SNPs revealed that *L. camara* in India exhibits a strong genetic structure. Interestingly, the structuring pattern does not exhibit a strong correlation with geography. In the structure analysis, individuals with similar flower colours clustered together regardless of their location of origin. The genetic distance between most of the individuals was low, indicating the absence of multiple species. A high inbreeding coefficient and a low proportion of heterozygous sites observed suggested that the strong structure could be due to self-fertilization. This was further confirmed by bagging experiments, which demonstrated that *L. camara* is self-compatible in India. Thus, we infer that *L. camara* exists as homozygous inbred lines formed by self-fertilization and that these inbred lines could be associated with distinct flower colours. Together, this would explain the correlation between flower colour and genetic structure, and the lack of geographic structure. These results refute the argument that *L. camara* is a species complex and emphasize the importance of the mating system in shaping the patterns of diversity in this invasive species. Our findings highlight a hitherto unknown role for mating systems in invasive species, furthering our understanding of evolution in invasive species.

## 1. Introduction

Invasive species are one of the major causes of ongoing biodiversity loss and extinctions in the Anthropocene (Linders et al., 2019; Mollot et al., 2017). Understanding the ecological and evolutionary underpinnings of invasion success is critical to controlling the emergence of new invasive species. Evolutionary and population genetic theory suggests that among other factors, genetic variation is critical to demographic success and range expansion (Ørsted et al., 2019). Populations that have undergone a demographic bottleneck are expected to have low genetic variation, which can negatively impact their ability to survive and adapt (Olazcuaga et al., 2023).

Invasive species typically undergo demographic bottlenecks during colonization (Alves et al., 2022). Low propagule pressure (the number of individuals introduced) leads to low effective population size. Consequently, invasive populations are established under conditions of limited genetic variation (Dlugosch & Parker, 2008; Simberloff, 2009). Despite the theoretical expectation of low genetic variation, invasive populations paradoxically demonstrate the ability to adapt, thrive and reproduce in large numbers in these novel habitats. This phenomenon is called the genetic paradox of biological invasions (Estoup et al., 2016). While extensively debated, some studies support the presence of such low genetic variation (Alves et al., 2022) while others contradict it (Bastien Lavergne & Molofsky, 2007; Zhang et al., 2021). The inherent complexities of human-mediated introduction like the choice of source population, number of introductions, artificial hybridizations etc. along with evolutionary forces make the patterns of genetic variation in invasive populations complicated (Roman & Darling, 2007). Several factors such as propagule pressure, hybridizations, gene flow, mating system etc. play an important role in determining the amount of genetic variation in invasive populations (Jaspers et al., 2021).

The introduction of many individuals either through single or multiple events can bring high variation into the novel environment (Bastien Lavergne & Molofsky, 2007). The source populations from which these individuals originate can influence the patterns of genetic diversity in new populations (Ryan et al., 2019). Gene flow between multiple invasive populations can homogenize genetic variation (Vallejo-Marín et al., 2021). Intraspecific hybridizations between genetically distinct individuals can lead to the formation of new genotypes with novel allelic combinations, while hybridization with native species can aid in rapid adaptation (Blair & Hufbauer, 2010; Mesgarana et al., 2016). Mating systems play an important role in determining the pattern of genetic diversity, especially in plant populations with extreme variations like self-fertilization and outbreeding (Hamrick et al., 1992; Ness et al., 2010; Schoen & Brown, 1991). Theoretically, cross-fertilizing species have high heterozygosity, and hence high genetic variation (Koelling et al., 2011).

Conversely, with self-fertilization heterozygosity decreases by 50% every generation (Hedrick, 2011). This leads to high genome-wide homozygosity and individuals exist as homozygous inbred lines (Guo et al., 2008; Pérez De La Vega & Garda, 1997). In self-fertilizing species, neutral diversity is low due to low effective population size (Foxe et al., 2009; Guo et al., 2008; Koelling et al., 2011), and it results in high genetic drift. The combination of low gene flow, high drift and high homozygosity leads to a strong genetic structure in self-fertilizing species compared to cross-fertilizing species (Edwards et al., 2019; Koelling et al., 2011; Mairal et al., 2023). While mating systems play an important role in determining genetic diversity, to the best of our knowledge, their role in invasive species has not been specifically investigated.

Here we present the genetic diversity patterns of the globally invasive plant species, *Lantana camara* (Lantana hereafter) in India. Lantana is a perennial shrub native to Central and South America. During the colonial era, this species was introduced into many countries across the globe, like India, Australia and South Africa as a garden plant (Kannan et al., 2013). Currently, it is one of the most important invasive species in India with a significant impact on the native ecosystems. On average, Lantana threatens almost 44 per cent of the forest areas in India (Mungi et al., 2020). In the Bandipur tiger reserve in southern India, Lantana occupies at least 40 per cent of the land area (Thekaekara et al., 2016). Unlike many invasive species, Lantana exhibits extensive phenotypic variation in India (Negi et al., 2019), with the most striking being the variation in flower colour. By 1900, approximately 16 species of the genus Lantana were reported in India, potentially suggesting multiple introductions. The taxonomy of the genus Lantana is complicated due to the presence of polyphyletic groups (Lu-Irving et al., 2021). In the earlier taxonomy, many flower colour variants of Lantana have been classified as distinct species (Sanders, 2006). However, genetic validation of this classification, focusing on different flower colour variants has not been done. It is possible that active hybridization to produce new hybrids took place in botanical gardens (Goyal & Sharma, 2015; Urban et al., 2011). Previous studies have hypothesized that invasive Lantana is a species complex, potentially formed from such hybridization events (Goyal & Sharma, 2015; Sanders, 2006). Variation in Lantana flower colour lends support to the species complex hypothesis. As a result, the invasive Lantana is sometimes referred to as *Lantana camara* sensu lato (Goyal & Sharma, 2015). Detailed genetic studies are important in checking this hypothesis and characterising different species of this genus.

Despite its status as one of the most pervasive invasive species globally, our understanding of its biology and genetics remains limited. It is further complicated by uncertainties regarding its species identity. Investigating the genetic diversity patterns of Lantana presents an opportunity to improve our understanding regarding the uncertainties about the species identity, as well as the genetic paradox of biological invasions. This can help in improving our understanding of the taxonomy of Lantana itself. A previous population genetic study using microsatellite markers suggests that Lantana populations in India are genetically homogeneous (Ray & Quader, 2014). However, as discussed earlier, its ecological and demographic success is very high along with the high phenotypic variation. We use thousands of genome-wide Single Nucleotide Polymorphism (SNP) markers and genetic simulations to investigate the patterns of genetic diversity, spatial genetic structure and the factors shaping these patterns in Lantana populations in India. We use India as an example of a large, ecologically diverse landmass where Lantana has been highly successful. More specifically, we investigate i) the genetic diversity and population structure of Lantana populations in India, and ii) the evolutionary forces shaping the genetic diversity patterns of Lantana.

## 2. Material and Methods

### 2.1 Study system

*Lantana camara* is considered to be one of the hundred most invasive species in the world (Lowe et al., 2000). It was introduced across the globe as a garden plant because of its strikingly attractive flowers. The earliest recorded presence of Lantana in India comes from the British botanical garden in Kolkata in 1807 (Kannan et al., 2013). Currently, it is widely distributed in India occupying almost every terrestrial habitat (Mungi et al., 2020). In India, Lantana exhibits extensive phenotypic variation with the most remarkable being the variation in the flower colour (Negi et al., 2019). Lantana produces umbellate inflorescence with younger flowers in the centre and older ones in the periphery. The colour of the flowers may change post-pollination, thus a single inflorescence can have flowers of different colours (Negi et al., 2019).

Flower colour variants were classified based on the colours of both young and mature flowers. For example, the ‘yellow–pink’ variant bears yellow flowers at the centre of the inflorescence that turn pink with age. In cases where flowers showed little or no colour change, variants were named according to their uniform colour, such as ‘orange’ (Fig. 1b). Yellow-pink, White-pink and Orange flower colour types are commonly observed in India (Fig. 1b). Flower colours like reddish-pink, yellow-orange and pink (Fig. 1b) are relatively uncommon. A single location can have multiple of these flower colour variants growing together next to each other.

**Figure 1.**
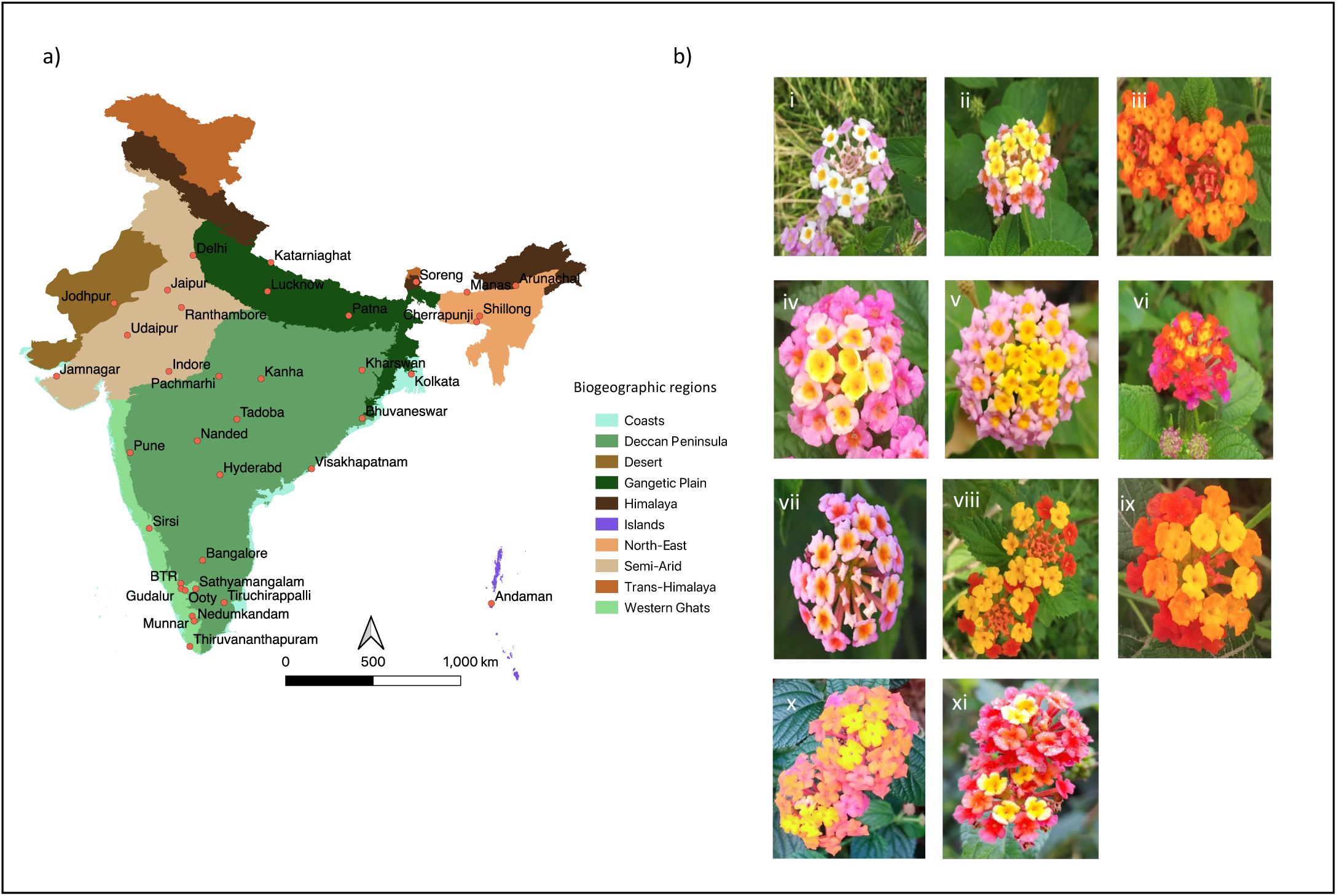
Sampling locations and flower colour variation in Lantana. (a) thirty-six locations were sampled across different biogeographic regions in India. (b) Flower colour variants used in the study. The names are given based on the variation within an inflorescence. i - White-pink, ii - Yellow-pink, iii - orange, iv - Yellow-pink-b, v - Yellow-pink-c, vi - Reddish-pink, vii - pink, viii - yellow-orange, ix - yellow-orange-b, x – Yellow-dark-pink a, xi - Yellow-dark-pink b.

### 2.2 Sampling locations

Lantana plant samples were collected between the years 2020 and 2023. Plants were collected from 36 locations across the country, representing various bioclimatic regions (Fig. 1a, Supplementary Table 1). Wild-growing plant saplings were collected and planted in the greenhouse at the National Centre for Biological Sciences, Bengaluru. From each location, a few plants that represented different flower colour types were selected randomly for the genomic analysis. We received silica gel-dried leaf samples from four locations which were also included in the study. Whenever possible, we have noted the flower colour during collection, or in the greenhouse. However, the exact flower colour of 59 plants that died before flowering is unknown.

### 2.3 DNA extraction and sequencing library preparation

Two to three fresh leaves were collected from each plant for DNA extraction. Leaves were crushed using liquid nitrogen in a mortar and pestle. A Qiagen DNeasy plant pro kit was used for the extractions, following the manufacturer’s instructions. DNA was quantified using a Qubit fluorometer. 199 μl Qubit buffer and 1 μl dye were mixed and 199 μl of this mix was used with 1μl DNA to check the concentrations.

Extracted DNA was stored at −20 degrees Celsius. Double-digest Restriction-site-associated DNA (ddRAD) libraries were prepared using the modified protocol by (Tyagi et al., 2024). Briefly, genomic DNA was digested using two restriction enzymes; SphI a six-base pair and MlucI a four-base pair cutter. Adapters specifically designed for SphI and MlucI are ligated to the cut ends. Unbound adapters were removed using magnetic bead-based cleaning, where 0.5X beads were used to remove fragments shorter than 50 base pairs. Individual specific index combinations were ligated to each sample using PCR and amplification was carried out. The PCR products were dual size selected for a target library size of 200 to 500 bp using magnetic beads. Library quality was evaluated using a TapeStation automated electrophoresis system. All the samples were pooled to get a 2nM library and the final library was sequenced in HiSeq2500 and NovaSeq6000 platforms.

### 2.4 Variant calling and filtering

Raw reads were demultiplexed and barcodes were trimmed using a custom script. Raw reads were quality-controlled using Trimmonatic - v.0.33.0 (Bolger et al., 2014). Reads with an average quality lower than 30 and reads shorter than 30 bp were excluded. Trimmed reads were aligned to a draft reference genome of *L. camara* (Joshi et al, 2022) using BWA-MEM (Li & Durbin, 2009) with default settings. We used Freebayes (Garrison & Marth, 2012) to call variants using the default settings, which were filtered using VCFtools (Danecek et al., 2011) to retain only high-quality SNPs. The filtering was done in the following order: biallelic SNPs were retained, SNPs with a minimum depth of 10 and a maximum depth of 500 were retained. Sites with a minimum genotyping quality of 30 and a minor allele count of at least three were kept. Individuals with more than 90% missing data were removed.

SNPs that are not in Hardy-Weinberg equilibrium were removed to minimize the influence of potential sequencing or genotyping artefacts. Finally, SNPs present in at least 90 per cent of the individuals were retained. The filtered VCF was used for all further analyses.

### 2.5 Population structure and genetic diversity estimations

We used a maximum likelihood based approach (ADMIXTURE) and a multivariate approach (Principal Component Analysis) to understand the genetic structuring of Lantana. Principal Component Analysis (PCA) was carried out using PLINK (Purcell et al., 2007). Genetic population structuring was investigated using ADMIXTURE (Alexander et al., 2009), with K values ranging from 2 to 17. We used Evanno’s delta K method (Evanno et al., 2005) implemented in the web server CLUMPAK (Kopelman et al., 2015) to find the optimum K. CLUMPAK was used to plot the admixture proportions as well. The structure plot was rearranged for easy visualization. Inbreeding coefficients and proportion of heterozygous sites were estimated using the R package Adegenet (v2.1.10) (Jombart, 2008). F_ST_ was estimated using Weir and Cockerham’s method implemented in VCFtools (Danecek et al., 2011). Adegenet (v2.1.10) was used to test for isolation by distance. Nei’s genetic distance was calculated using the StAMPP package (Pembleton et al., 2013) in R. A phylogenetic network was generated using the r package StAMPP and the SplitsTree4 (Huson, 1997). We used a MANOVA to test the association between flower colour and genetic structure, based on assignment proportions at K = 11. We also plotted the assignment proportions of the major flower colours to visualize the relationship between genetic structuring and flower colour.

### 2.6 Demographic history analysis

We used demographic history simulations in fastsimcoal2 (Excoffier et al., 2021) to test whether the divergence among flower colour variants of Lantana predated its introduction. Three flower colour morphs from a single population in Bhubaneswar, eastern India were analysed in pairwise comparisons: yellow–pink (4 individuals), white–pink (4 individuals), and orange (3 individuals). Divergence was assessed between the white–pink and orange morphs, and between the white–pink and yellow–pink morphs. VCF files were generated for these individuals and the input site frequency spectrum (SFS) was generated with easySFS (https://github.com/isaacovercast/easySFS). Ten demographic scenarios were tested, broadly falling into two categories: (i) divergence predating the species’ introduction and (ii) divergence occurring after introduction, with or without gene flow at different stages of demographic history (Supplementary Fig. 5). Based on historical records, the bottleneck was hypothesised to have started ∼350 to 450 years ago and ended ∼150 to 250 years ago (including introduction to Europe) (Kannan et al., 2013). These values were used as parameter ranges in the simulations. A generation time of two years was considered. Simulations were run with parameters -n 50000 -m -M −l 30 -L 60 -E 1000. The best-fitting model was selected using the Akaike Information Criterion (AIC), with the lowest AIC indicating the most likely scenario. Ten replicates of the selected model were run to estimate confidence intervals for the parameter values.

### 2.7 Bagging experiment

Lantana is widely considered to be a cross-fertilizing species with insect-mediated pollination. To investigate the mode of pollination in Lantana, we conducted a bagging experiment. Mature but unopened inflorescences were enclosed in fine mesh bags to exclude pollinators, while inflorescences at the same developmental stage were tagged as unbagged controls and left open for natural pollination. The experiment was performed on wild growing plants of the yellow-pink flower colour on the GKVK campus, Bangalore. To prevent insect entry, the mouth of the bags were smeared with a sticky barrier. In total, 33 inflorescences were bagged and 33 were tagged as open-pollinated controls. After flowering and fruit maturation, seed set was recorded for both treatments. A t-test was used to assess significant differences between treatments.

### 2.8 Genetic simulations

We used SLiM (Haller & Messer, 2019), a forward-in-time, individual-based simulation program to understand how mating systems influence the pattern of genetic diversity in invasive plant species. Populations analogous to both native and invasive populations were simulated with different mating systems. All the individuals in the simulations were diploid hermaphrodites, with 10 chromosomes each with 1,00,000 bps. The mutation rate and the recombination rate were considered to be 10^-8^ mutations per site per generation and 10^-7^ recombination per base pairs per generation respectively (Kovalchuk et al., 2000; Waneka et al., 2024), which is the general mutation and recombination rate for plants. Generations were assumed to be non-overlapping. Scenarios with different mating systems having varying levels of self-fertilization (zero - complete crossing, 25, 50, 75 - mixed-mating and 100 - complete selfing) were simulated. A schematic representation of the demographic changes during the simulation is shown in Figure 2. The simulation started with a single population of 100 genetically identical individuals, which grew exponentially (growth rate = 1.05) until they reached a carrying capacity of 10,000 individuals. This carrying capacity was decided, to reduce the computational complexity. In the burn-in phase, the individuals evolve under these conditions for 20,000 years/generations. In the 20,000^th^ year, two new populations were created by individuals migrating from the initial simulated population (assumed to be the native population). The number of founding individuals (in this case propagule pressure) was either 10, 100 or 1000.

**Figure 2.**
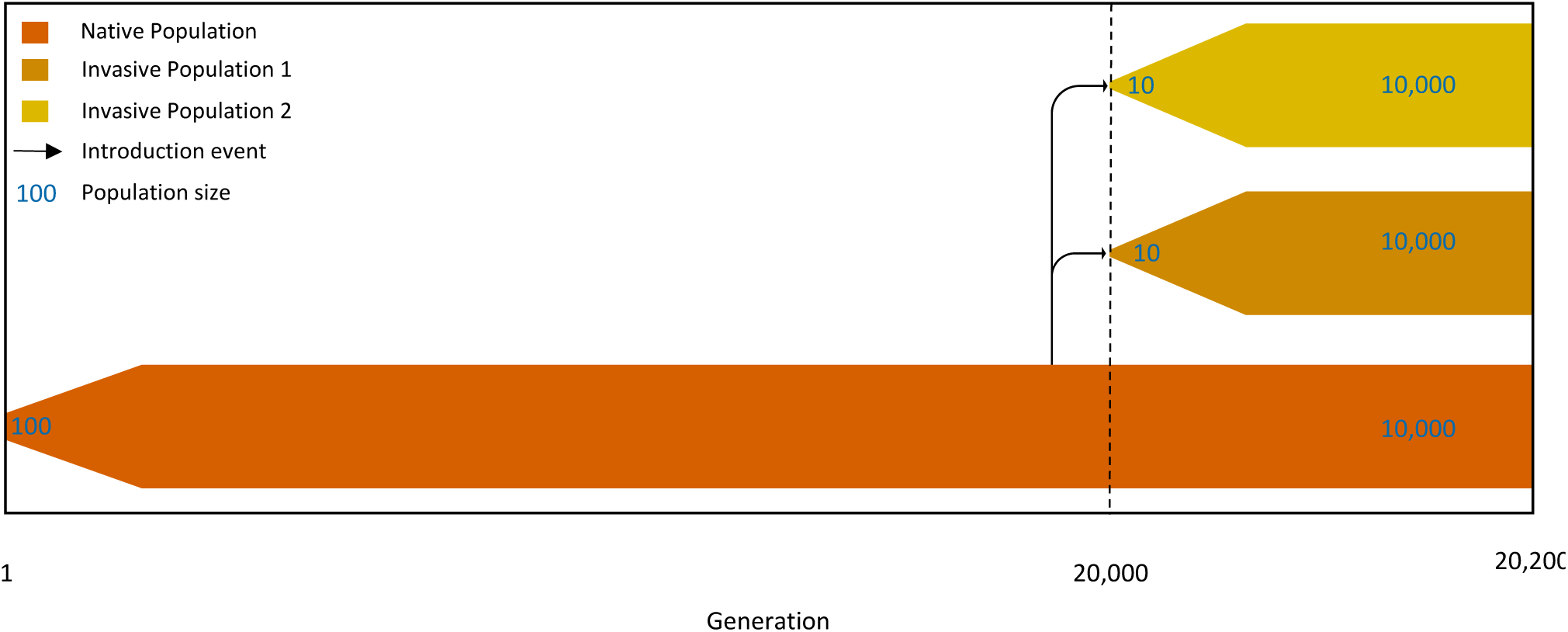
Schematics of the populations modelled in the SLiM simulations. The simulations begin with a single population of 100 individuals, which expands exponentially until reaching a carrying capacity of 10,000 individuals. At generation 20,000 two new invasive populations are founded by 10 individuals each. All three populations then evolve independently for an additional 200 generations.

These populations grew exponentially to a carrying capacity of 10,000 with the same growth rate. The population evolved till the year 20,200 and thus 200 generations after invasion is simulated. The population level heterozygosity, nucleotide diversity and F_ST_ between populations were measured at the end of the 20^th^ and 200^th^ generations. For randomly selected 50 individuals from each population, VCF files with SNP information were created at the 20,200^th^ generation.

## 3 Results

### 3.1 Invasive Lantana populations in India show strong genetic structure

Using ddRAD sequencing, we obtained 19,008 SNP markers for 359 Lantana individuals from different regions across India. This SNP dataset was utilised to assess the genetic differentiation in Lantana through four approaches: ADMIXTURE, Principal Component Analyses, phylogenetic network reconstruction and estimation of fixation index. Structure analyses using the ADMIXTURE package revealed strong genetic structure in the Indian Lantana populations (Fig. 3a). The optimum number of Hardy-Weinberg populations was between four and eleven (Supplementary Fig. 1).

**Figure 3.**
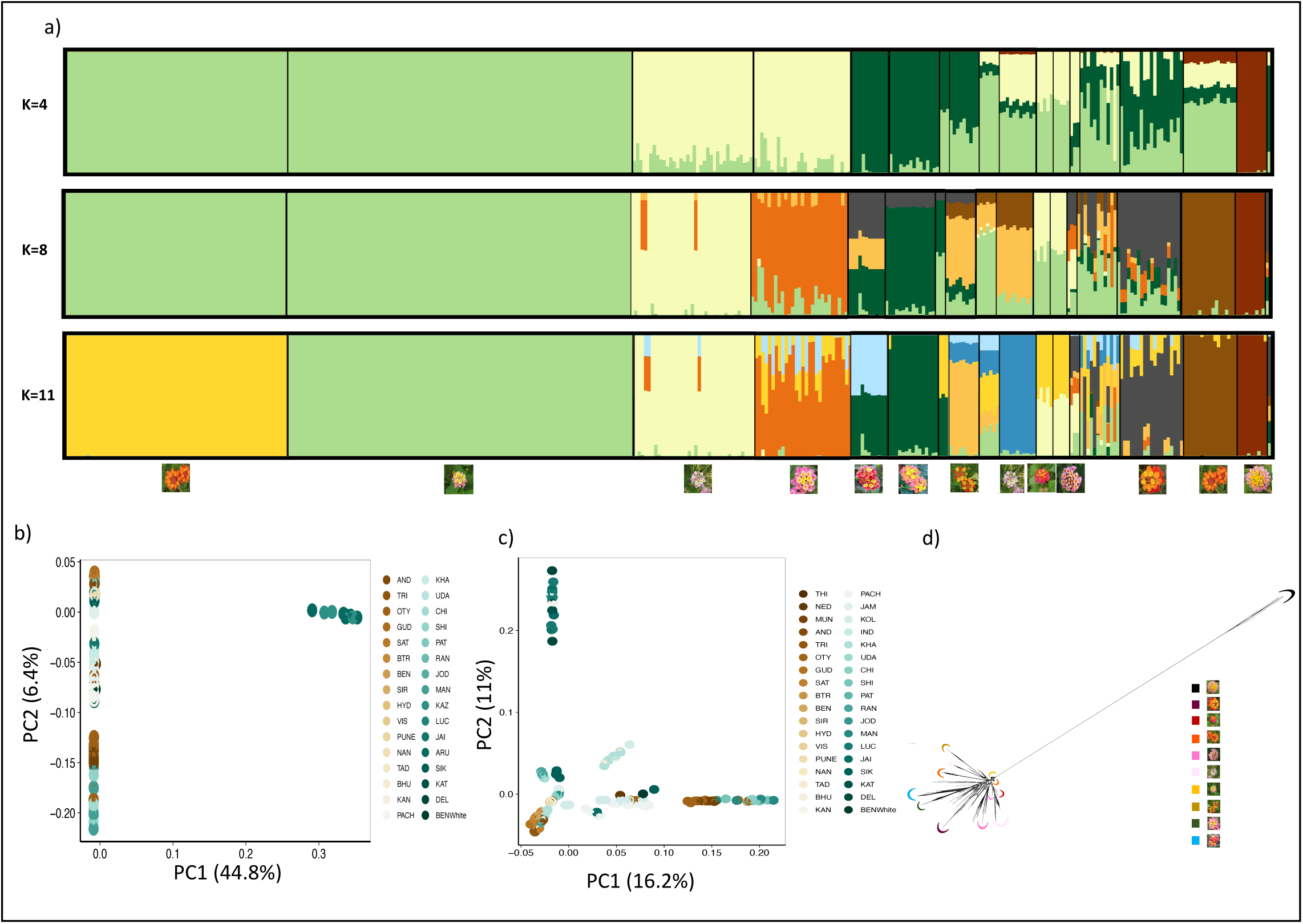
Population structure of Lantana in India. a) Genetic ancestry assignment using ADMIXTURE programme. Ancestry fractions for K = 4, K = 8 and K = 11 are given with clusters arranged by flower colour variants b) PCA of all the 359 samples c) PCA excluding nine samples from northeast India. Individuals are coloured based on their geographic origin from north (green) to south (brown) of India d) Phylogenetic network showing different lineages associated with flower colour, lineages are coloured based on the flower colour.

At K=4, which had the highest support for the optimum number of clusters, the geographic origin did not dictate the genetic structuring. Despite belonging to the same geographic location, individuals were assigned across different genetic clusters. At K=3, nine individuals from North-Eastern India clustered together and maintained a distinctive identity at higher K as well. Many individuals were admixed. K=8 and 11 had the second and third highest log likelihood values for the optimum K.

The principal component analysis results corroborated the findings from the admixture analyses (Fig. 3b and c). In the PCA encompassing all 359 samples (Fig.3b), PC 1 explained 44.8% of the variation and PC 2 explained 6.4% of the total variation. Nine samples from North-eastern India formed a distinct cluster separate from the rest of the samples in the PC1 axis (Fig. 3b). To explore the structuring pattern within the remaining samples, a separate analysis excluding these nine individuals was carried out (Fig. 3c). In this analysis, PC 1 explained 16.2% of the variation and PC 2 explained 11% of the variation. In the PCA, samples were coloured based on the geographic location from south to north of India (brown to green respectively). Individuals from geographically distant locations were clustering together while those from the same geographic area were assigned to different clusters in many locations, reinforcing the absence of a strong geography-based pattern. A flower colour-based clustering pattern was evident in the PCA with samples colour-coded according to the flower colour (Supplementary Fig. 2).

In the phylogenetic network generated using SplitsTree (Fig. 3d), the nine samples for the north eastern India clustered separately. The isolation by distance analysis using the Mantel test (Supplementary Fig. 4) revealed a weak correlation between genetic distance and geographic distance (Mantel r = 0.039, p-value = 0.075), indicating that geography is not the major factor influencing the structuring pattern. The fixation index (F_ST_) between most of the population was low (Supplementary Table 2). Arunachal Pradesh (in north-eastern India) samples showed high F_ST_ with many populations (0.46 to 0.54) (Supplementary Table 2).

### 3.2 Genetic population structuring in Lantana is associated with flower colour

In the Admixture analysis, the clusters closely aligned with distinct flower colour variants of Lantana. At K=4, yellow-pink and some of the orange-flowered Lantana clustered together, while a separate cluster comprised pink and some of the white-pink individuals (Fig. 3a). A third cluster was formed by dark-pink-coloured plants from Ooty (Southern India) and North-east India. Nine individuals from northeast India formed a fourth cluster. Other flower colour variants were admixed.

Interestingly, at K=11, a pronounced flower colour-based clustering pattern emerged, with distinct clusters formed by yellow-pink, pink, reddish-pink, dark pink and yellow-orange individuals, while both orange and white-pink Lantana each formed two separate clusters. In the PCA, the clustering pattern often mirrored the flower colour variants (Supplementary Fig. 2). Yellow-pink colour variant formed a tight cluster.

One of the orange colour variants also showed a similar trend. In the phylogenetic network generated using SplitsTree, (Fig. 3d) most of the individuals were grouped according to their flower colour. The most common flower colour in the country, yellow-pink, formed a single cluster in admixture analysis and in the phylogenetic network. Yellow-pink and orange individuals showed short branch lengths in the phylogenetic network. The MANOVA revealed a strong correlation (Pillai’s trace = 3.07, approx. *F* (50, 1270) = 40.27, *p* < 2.2 × 10^-16^) between flower colour and genetic structure (K = 11), with individuals sharing similar flower colours assigned together in a cluster (Supplementary Table 5). The visual representation of flower colour allocations across genetic clusters also illustrates the association between flower colour and genetic structuring (Supplementary Fig. 7).

### 3.3 High Inbreeding and low genetic diversity characterize Lantana populations

The Nei’s genetic distance among most of the Lantana individuals across the country was very low, indicating the absence of genetically distinct species, contrary to the earlier hypotheses (Fig. 4a). Despite this overall low genetic distance, strong differentiation was often observed within populations, as revealed by the genetic differentiation analysis. Notably, high genetic distance was detected only for the nine individuals from Northeast India relative to the rest of the samples. This pattern was consistent with the phylogenetic network, where these nine individuals formed a distinct group with long branch lengths (Fig. 3d). The inbreeding coefficient was extremely high for many individuals (Fig. 4b). Nearly all the individuals exhibited a very low proportion of heterozygous sites, with less than 10% heterozygous sites (Fig. 4c). However, the nine individuals from North-East India showed the highest proportion of heterozygous sites (>0.3). Taken together with the genetic differentiation analysis, these results indicate a predominantly self-fertilizing mating system in Lantana.

**Figure 4.**
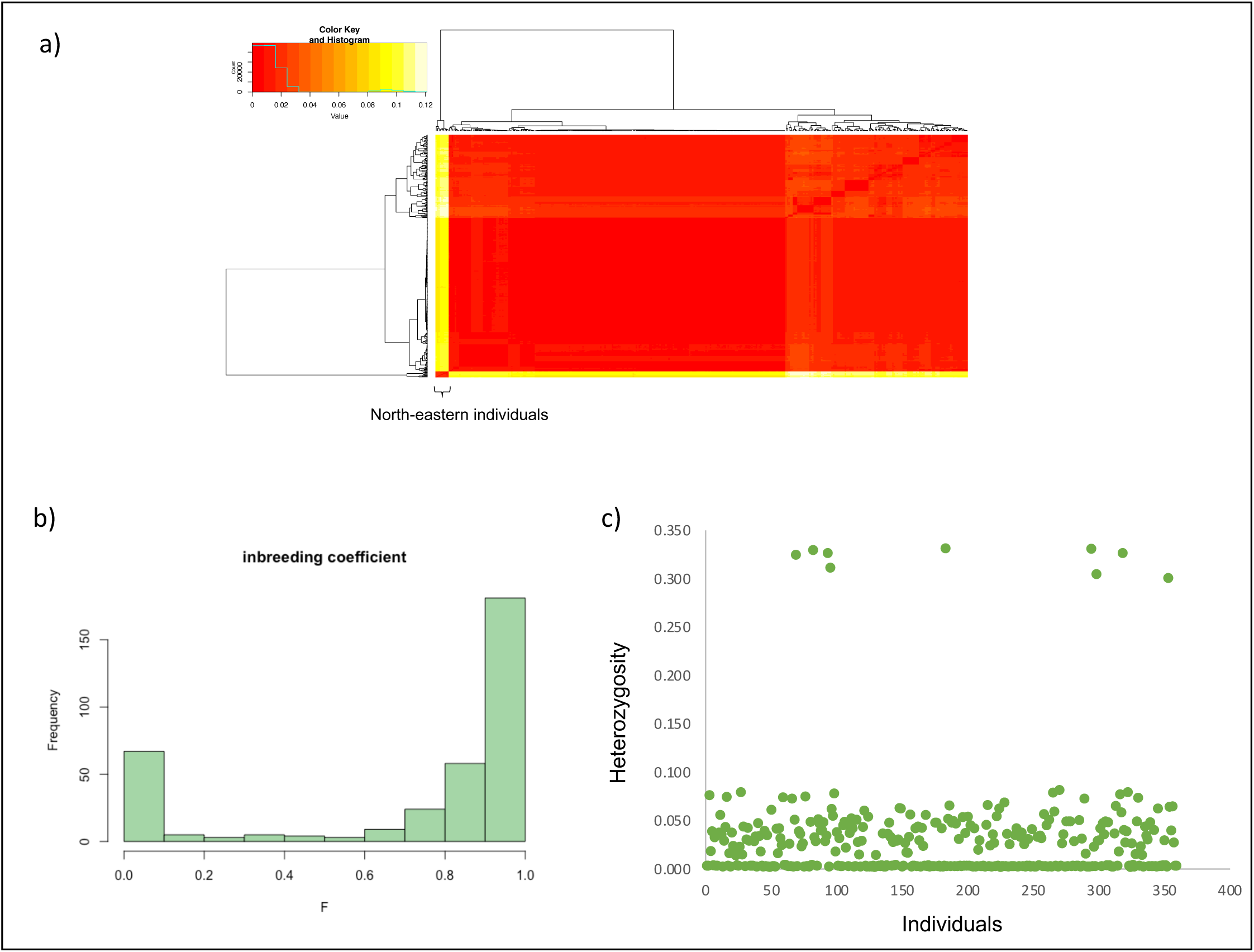
Genomic signatures indicating high inbreeding in Lantana a) Nei’s genetic distance among all the individuals. Nine samples form northeast Indian are genetically most distinct b) Inbreeding coefficient (F) c) Proportion of heterozygous sites per individual.

To test whether Lantana is self-compatible in India, we conducted a bagging experiment. A total of 33 inflorescences were enclosed in mesh bags, while an equal number were left open to natural pollination. All open-pollinated inflorescences set seed, and all but one of the bagged inflorescences also produced seeds. In total, 370 seeds were obtained from bagged inflorescences and 402 seeds from open-pollinated ones. Seed set did not differ significantly between treatments (t = 0.76, df = 64, p = 0.45), indicating self-compatibility and a predominantly self-fertilizing mating system (Fig. 5b). These results confirm that Lantana is self-compatible in India.

**Figure 5.**
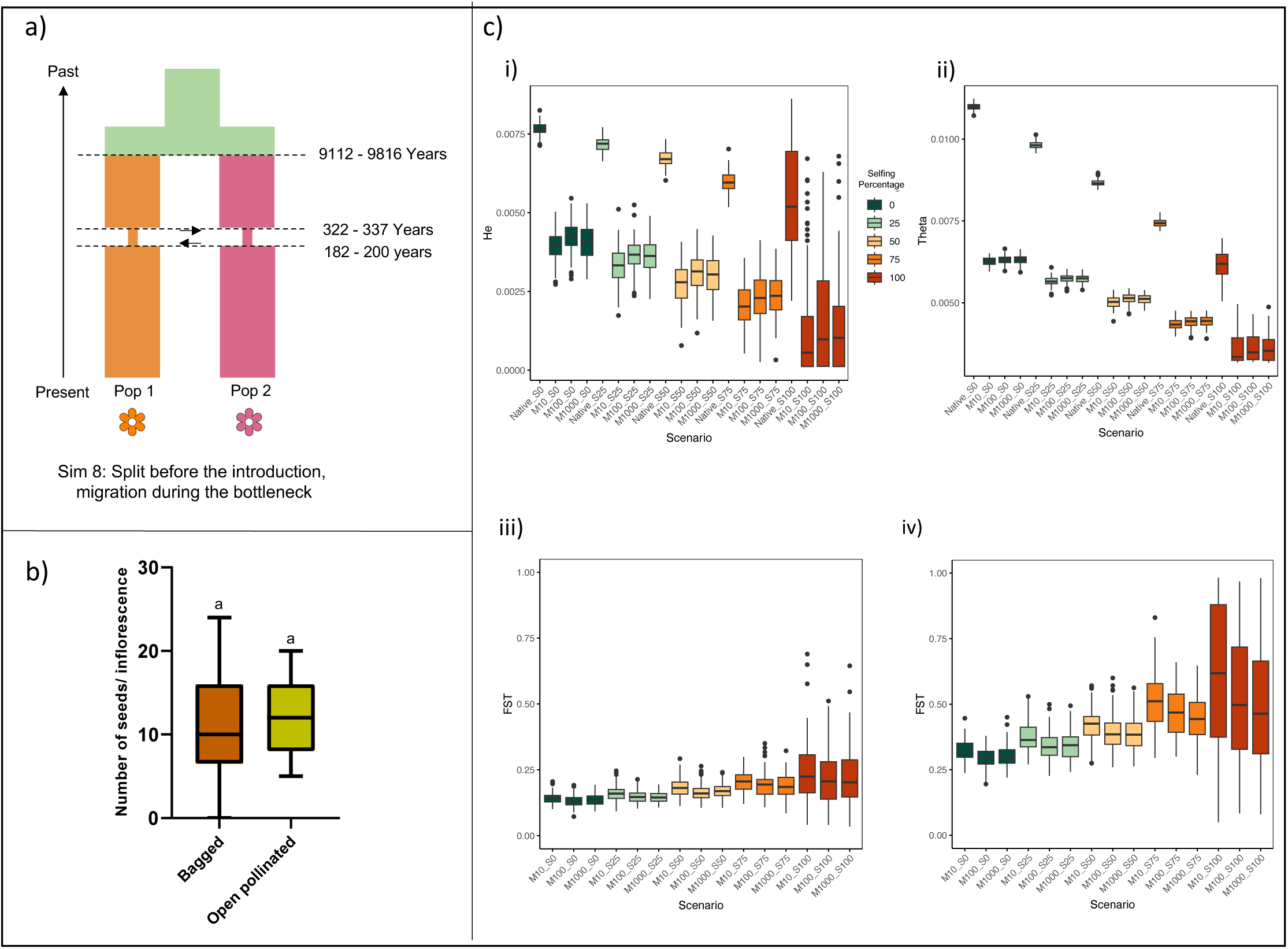
Evidence supporting the predominance of self-fertilization in Lantana (a) Best-supported demographic model inferred using fastsimcoal2. Pop1 and Pop2 represent distinct flower colour lineages, with arrows indicating inferred gene flow (b) Graph showing results of the bagging experiment (c) Individual-based genetic simulation results showing putative patterns of genetic diversity in invasive plants under different assumed mating systems and propagule pressure (i) Heterozygosity (ii) and nucleotide diversity (iii) under different mating systems in ‘native’ and ‘invasive’ populations founded by 10, 100 and 1000 individuals c) F_ST_ between ‘native’ and ‘invasive populations’ under different simulated mating systems and propagule pressure. (iv) F_ST_ between two simulated ‘invasive populations’ under different mating systems and propagule pressure.

To determine whether the divergence between lineages occurred before or after the introduction, we inferred demographic history using fastsimcoal2. We simulated demographic history scenarios with divergence predating the introduction or divergence occurring after the introduction, incorporating gene flow at different stages. Among the ten alternative models tested (Supplementary Fig. 5), *Sim 8* had the lowest AIC and was selected as the best fit model (Fig. 5a, Supplementary Table 3, Supplementary Fig. 6) for both white-pink yellow-pink divergence and white-pink orange divergence. This model indicates that the divergence between flower colour morphs predates the introduction, with a bottleneck beginning between 322 to 337 years ago and ending between 182 to 200 years ago (Supplementary Table 4) for white-pink and orange. Similar results were found with slight variation for yellow-pink and white-pink as well (Supplementary Table 4, Supplementary Fig. 6). It also indicates gene flow during the period of the bottleneck or introduction, consistent with earlier proposals of artificial hybridization in botanical gardens. Collectively, these results suggest that multiple pre-existing lineages that evolved within the native range were introduced into India.

### 3.4 Genetic diversity in invasive plant populations is strongly influenced by the mating system

To understand the influence of the mating system on genetic diversity in invasive plant species, we have carried out an individual-based genetic simulation using SLiM. These simulations confirmed that the mating system strongly impacts standing genetic variation within an invasive plant species. Simulations of cross-fertilisation resulted in higher heterozygosity and nucleotide diversity than those of self-fertilization (Fig. 5 c-i and c-ii). This was consistent across various propagule pressures (10, 100 and 1000) simulated. The heterozygosity and nucleotide diversity were low in the simulated invasive populations compared to the simulated native population (Fig. 5 c-i and c-ii). An increase in the propagule pressure from 10 to 1000 did not substantially increase the genetic diversity. Under the same propagule pressure, simulations of cross-fertilization gave rise to invasive populations with high genetic diversity compared to scenarios with self-fertilization. The variance in heterozygosity and nucleotide diversity was high in the self-fertilizing simulation scenarios.

Mating systems had an impact on the differentiation of simulated native and invasive populations. F_ST_ between simulated native and invasive populations revealed that the F_ST_ value was lowest under complete cross-fertilization (Fig. 5 c-iii). The differentiation increased as the percentage of self-fertilization increased. Under simulations of complete self-fertilization, the variance was high and F_ST_ ranged from approximately 0.03 to 0.69. Whereas, in complete cross-fertilization, it ranged from 0.07 to 0.2. Similarly, in simulations of cross-fertilization, the differentiation between simulated invasive populations was low compared to self-fertilisation scenarios (Fig. 5 c-iv). The differentiation increased as the allowed selfing percentage increased. F_ST_ ranged from 0.05 to 0.98 in self-fertilizing simulation scenarios. Within each category of selfing percentage, an increase in the propagule pressure reduced the differentiation.

### 3.5 Self-fertilization leads to strong structuring within the simulated populations

PCA using SNPs from individuals of all three populations revealed a strong genetic structure in self-fertilization scenarios (Supplementary Fig. 3). Whereas, in cross-fertilization scenarios, there was a weak structure (Supplementary Fig. 3).

Under selfing, individuals from the same population were assigned to distinct genetic clusters, and within a cluster, individuals from multiple populations were present, similar to what we have observed in Lantana populations.

## 4 Discussion

Biological invasions pose a serious global challenge, with millions of dollars spent annually on their control (IPBES, 2023). Yet, it is often difficult to control the emergence of new invasive species and their spread due to the lack of understanding of the invasion process and the factors contributing to invasion success. Critical to understanding invasion success is an understanding of standing genetic variation and population differentiation. We used genomic techniques to understand patterns of genetic diversity and differentiation in invasive Lantana populations in India. Our findings revealed that Lantana populations in India are genetically structured, but the structuring pattern was not strongly correlated with the geography. In the Admixture analysis, the genetic structure aligned with flower colour variation. Notably, many individuals exhibited a high inbreeding coefficient and a very low proportion of heterozygous sites. Genetic distance indicated that lineages are relatively similar. We suggest that the strong genetic structure despite the low genetic distance is because of the increased homozygosity due to self-fertilization.

The bagging experiment confirmed that Lantana is self-compatible, supporting the results from genomic analyses. Demographic simulations further suggest that these lineages or inbred lines likely originated prior to the introductions. Through genetic simulations, we show that the mating system can impact genetic diversity and the genetic structure of simulated, putatively invasive species.

### 4.1 Population structure

Our investigation into the genetic structure of Lantana using both Admixture and principal component analysis revealed a strong population structure. Individuals from the same geographic location were assigned to different genetic clusters, contrary to a previous study using microsatellite markers which suggested genetic homogeneity in Lantana populations in India (Ray & Quader, 2014). SNP markers are numerous in the genome, and as a result, have higher statistical power than microsatellite markers (Zimmerman et al., 2020). We have used 19,008 SNP markers and thus we expected population structure to be better resolved in our study.

In PCA, admixture, and phylogenetic network nine individuals from the northeast of India were genetically distinct compared to the rest of the samples. These samples exhibited high heterozygosity compared to other plants, which could explain their genetic distinctiveness. These plants also had noticeable phenotypic differences, such as a lack of thorns and dense trichomes on leaves. Further detailed analyses of these plants are required to better understand the basis of such a large genetic difference.

Many invasive species show strong genetic structure in the invaded ranges (Wang & Chen, 2017) often as a result of multiple introductions followed by limited gene flow between populations (Bastien Lavergne & Molofsky, 2007). However, in many such studies, a geography-based population structuring pattern is observed (Le Cam et al., 2020). In contrast, Lantana shows strong genetic structure, with weak correlation with the geographic distance. These genetic patterns are consistent with the expectation of multiple introductions, as suggested by historical records on Lantana (Kannan et al., 2013). As expected with multiple introductions, we find multiple genetic clusters across populations and accompanied by considerable phenotypic diversity especially in flower colours of Lantana in India. However, our findings are unique because, despite the presence of multiple genetic clusters, there is no evidence of a strong geographic structuring.

### 4.2 Self-pollination leading to high homozygosity in Lantana

Most Lantana individuals in India exhibit a high inbreeding coefficient and a low proportion of heterozygous sites. In plants, such patterns are typically associated with self-fertilization (Hedrick, 2011). Although these patterns are not exclusive to self-fertilization, when considered alongside the genetic differentiation results, they strongly suggest that Lantana in India is predominantly self-fertilizing and persists as homozygous inbred lines. The bagging experiment provided further evidence of self-compatibility, supporting the conclusion that Lantana primarily reproduces through self-fertilization. In many natural populations of plants, the mating system is shown to strongly influence the patterns of genetic diversity (Bomblies et al., 2010). Mating systems in plants range from uniparental reproduction or self-fertilization to obligate cross-fertilization or outcrossing (Charlesworth, 2006). Approximately 10-15% of seed plants predominantly self-fertilize (Goodwillie et al., 2005). Many species exhibit mixed mating systems, including a combination of both selfing and outcrossing (Whitehead et al., 2018). Mating systems influence the effective population size and thus play a crucial role in shaping genetic diversity within a species (Laenen et al., 2018).

It is reported that in its native range, Lantana primarily relies on cross-fertilization for reproduction (Barrows, 1976). However, in the introduced range, Lantana is self-compatible (Goulson & Derwent, 2004; Ram & Mathur, 1984). During colonization and invasion, plant species can shift mating systems from cross-fertilization to self-fertilization (Petanidou et al., 2012; Willi et al., 2022). However, the demographic history simulations indicate that the divergence between lineages predated the introduction, suggesting that self-fertilization may already have been present in the native populations. A detailed comparative study of mating systems in both the native and invaded ranges is therefore required to confirm whether such a shift occurred in Lantana. Self-fertilization is shown to increase the genome-wide homozygosity and inbreeding coefficient (Barragan et al., 2024). The strong genetic structure even with low genetic differences can be attributed to the high homozygosity of the inbred lines formed through selfing (Bomblies et al., 2010). The presence of multiple genetic groups in a single location is mainly due to the presence of such inbred lines. This is supported by studies like Bhattacharya et al., 2022.

Selfing acts as a barrier to gene flow between these inbred lines, thereby maintaining their genetic identity. However, the genetic distance between these lines suggests that they are not genetically very different. The presence of admixed individuals in many populations indicates the occurrence of cross-fertilization as well. These occasional crossing events can help in generating new recombinant inbred lines.

### 4.3 Association of flower colour with genetic structure

Our study revealed an association between genetic clustering and flower colour variation. Different flower colour variants were assigned to distinct genetic clusters despite being from the same geographic location. This indicates the lack of gene flow between flower colour variants which grow physically close to each other in many locations. This correlation between flower colour and genetic structuring underscores the influence of the mating system. Here, self-fertilization acts as a barrier to gene flow and different inbred lines are associated with different flower colours. Thus, we think that the identity of these lines is maintained by self-fertilization. Historically, Lantana species were primarily defined mainly based on the flower colours and many such variants were considered as different species (Sanders, 2006). Our study suggests that these are within species variation and the overall phenotypic similarity within a flower colour variant is maintained by self-fertilization. We think that the correlation between flower colour and genetic structure may result from the formation of inbred lines. Further genetic analysis is required to understand the genetic basis of these flower colour variations.

### 4.4 Is Lantana a species complex?

The findings from the molecular analysis of Lantana challenge the notion of invasive Lantana being a species complex. While genetic structuring was evident, it predominantly arose from the prevalence of highly homozygous inbred lines rather than from the presence of genetically different species. Most lines exhibited low genetic distance between each other, indicating the high genetic similarity between flower colour variants. The high phenotypic variation observed could be the result of the introduction of multiple inbred lines to the subcontinent or the emergence of novel inbred lines through hybridization post-introduction. Our demographic history simulation suggested the introduction of genetically distinct lineages. The variations observed could be within species phenotypic variations typical to many plant species. Through citizen science platforms like iNaturalist, it is evident that Lantana in its native range also exhibits similar flower colour variation. However, genomic analysis of related species of Lantana can help in understanding the chances of any past introgressions.

## 5 Conclusions

Studying genetic diversity patterns in invasive species is crucial for understanding their evolution. We explored genetic diversity patterns of the global invasive species *L. camara*. Our results reveal that lantana populations in India are strongly structured with genetic differences between flower colour variants. The low genetic distance observed between most plants negates the chances of lantana being a species complex. The presence of strong genetic structure even in the absence of high genetic distance is attributed to self-fertilization and the formation of putative inbred lines. This is supported by the low heterozygosity, high inbreeding coefficient and the bagging experiment. In predominantly self-fertilizing species, individuals exist as inbred lines, which restricts gene flow and allows these inbred lines to maintain their genetic identity. We suggest that the correlation between flower colour and genetic structure is due to the association of specific flower colours to specific inbred lines.

Uniquely, our study highlights the significance of the mating system in shaping the genetic diversity pattern in invasive plant species. Our results also highlight the importance of multiple introductions and the introduction of genetically different variants in contributing to high genetic structuring within invasive populations. Further research on other invasive species will reveal how general these patterns might be.

## Acknowledgements

We are thankful to Keval Palya, Mamta M., Mayuresh Gangal, and Rajat Rastogi for their help with sampling and laboratory experiments. We are thankful to Divyashree Rana, Kritagnya Vadar, Nishma Dahal, Rachana Rao, Tikily Tayeng, Tista Ghosh and Vinay Sagar for aiding in the sample collection. We thank Mayuresh Gangal, Divyashree Rana, Abhinav Tyagi, Anubhab Khan, Vinay Sagar and Arpitha Jayanth for their valuable suggestions. We thank the forest departments of Karnataka (No: PCCF (WL) /E2/CR-52/2019-20), Tamil Nadu (No: WL(A)/52852/2019) and Madhya Pradesh (MP permit no:835, dated: 30-01-2020) for providing necessary permits. PP was supported by NCBS/TIFR (Department of Atomic Energy). This work was supported under DBT project No-BT/PR29251/FCB/125/18/2018 and the NCBS-TIFR internal plan fund that was awarded to UR. The NCBS data cluster used is supported under project 12-R&D-TFR-5.04-0900, Department of Atomic Energy, Government of India. For institutional support, we thank the National Centre for Biological Sciences (NCBS). We thank the Next Generation Genomics facility at NCBS, for helping with the sequencing.

## 6 Author contributions

Conceptualization: P.P., U. R, and R. G., Data collection: P.P., Laboratory work: P. P., Data analysis: P.P, Project administration: P.P., U.R., and R. G., Funding acquisition: U. R, and R. G., Writing original draft: P.P., Writing – Review and editing: U.R., Supervision: U.R.

## 7 Data availability statement

Accession numbers for the raw ddRAD sequencing data and the GitHub link for analysis codes will be provided upon acceptance of the manuscript.

## Supplementary material

**Figure 1.**
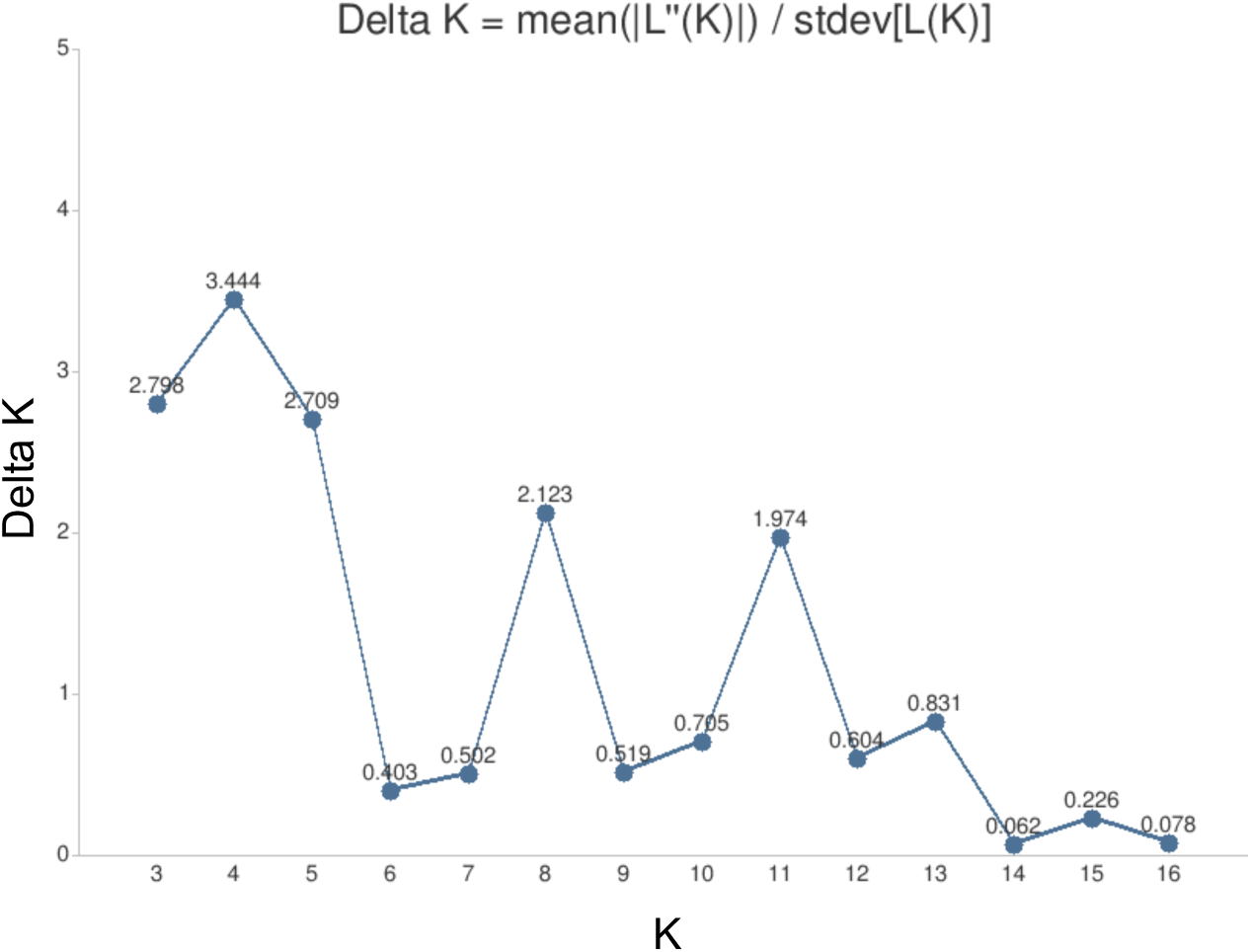
Optimum number of clusters using Evanno’s delta K method.

**Figure 2.**
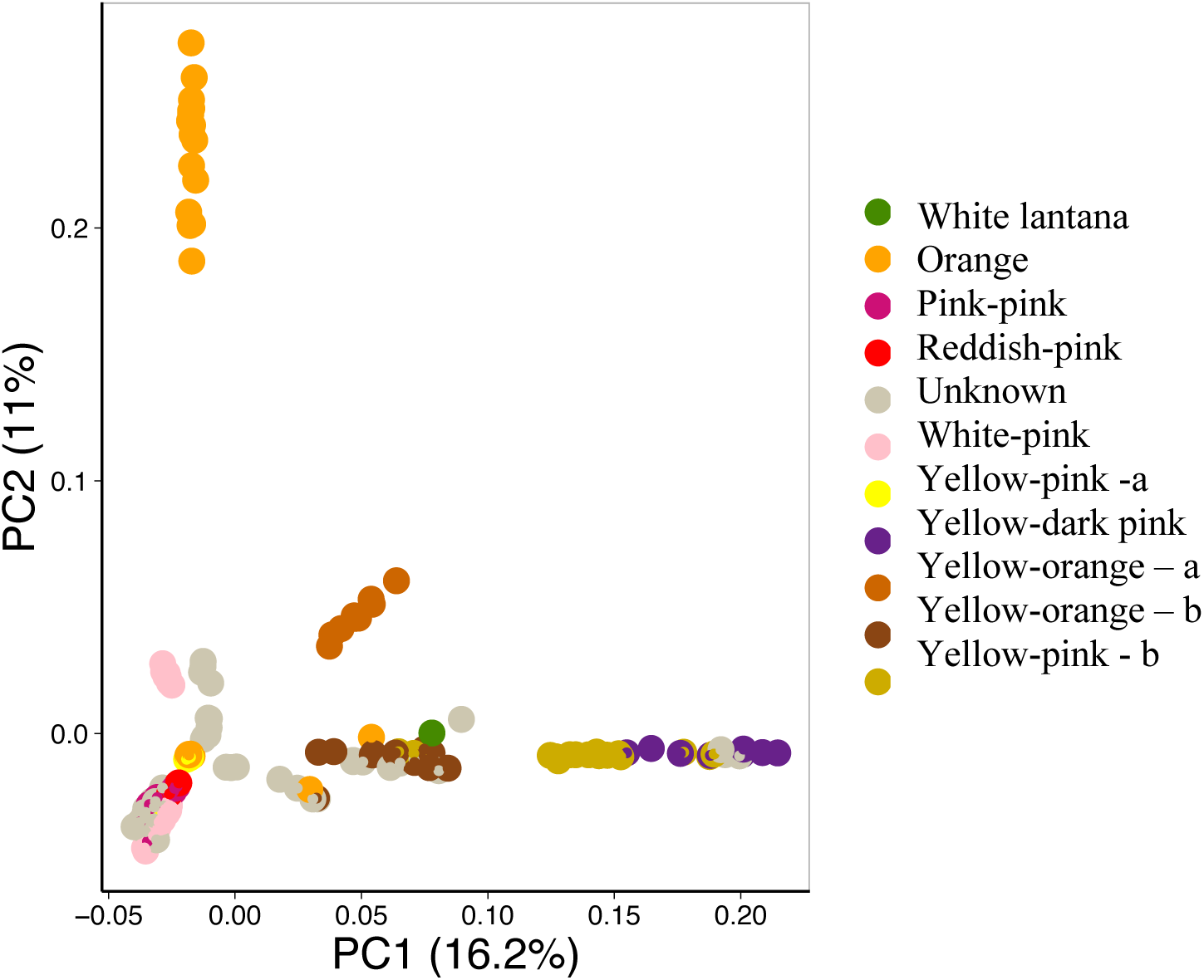
PCA based on flower colour of Lantana individuals in India. Individuals with similar flower colours are coloured similarly in the plot. The flower colour of some of the samples was unknown.

**Figure 3.**
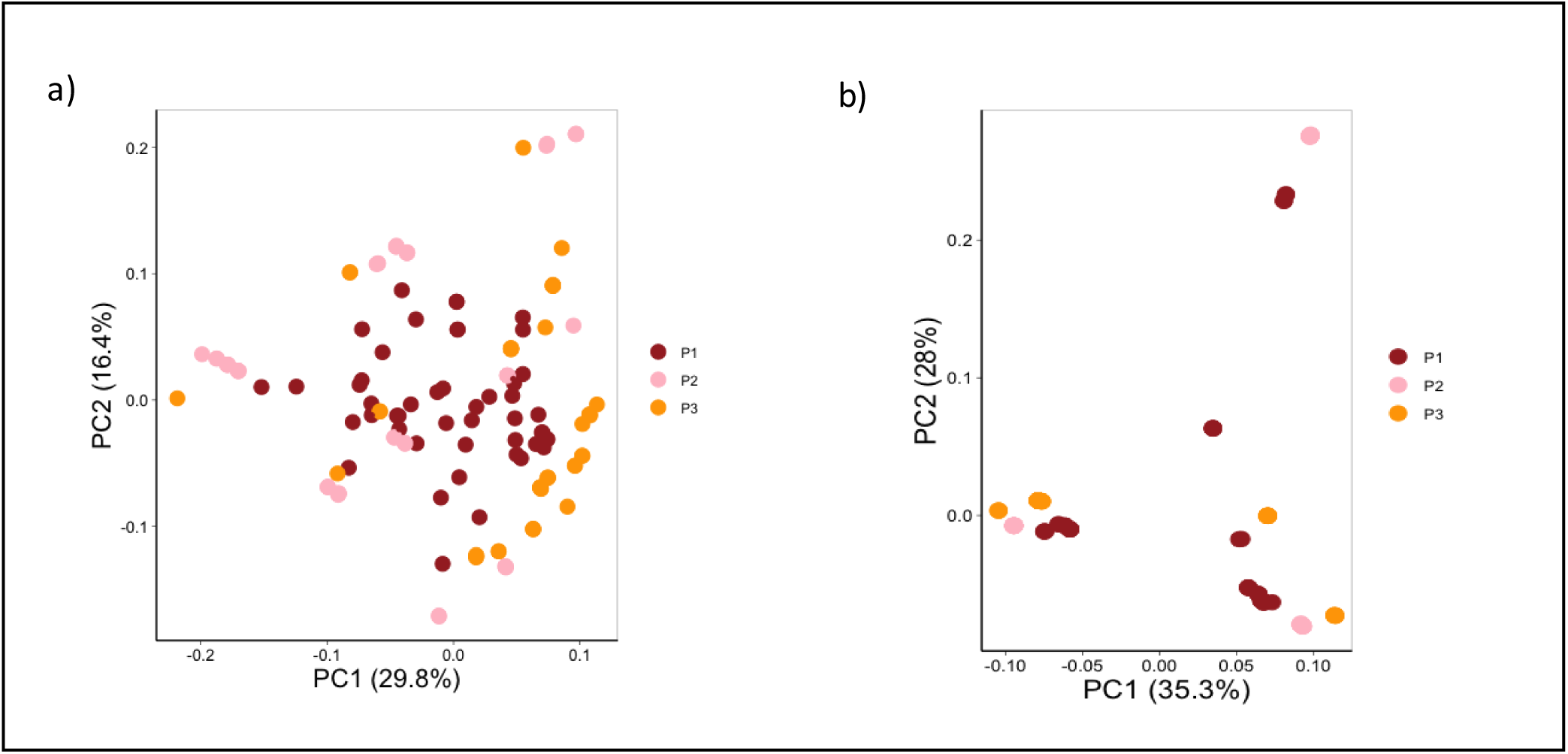
PCA of 50 randomly selected individuals from the simulation scenarios. P1 is the simulated native population and P2 and P3 are the simulated invasive populations. a) PCA of cross-fertilizing individuals simulated b) PCA of self-fertilizing individuals simulated

**Figure 4.**
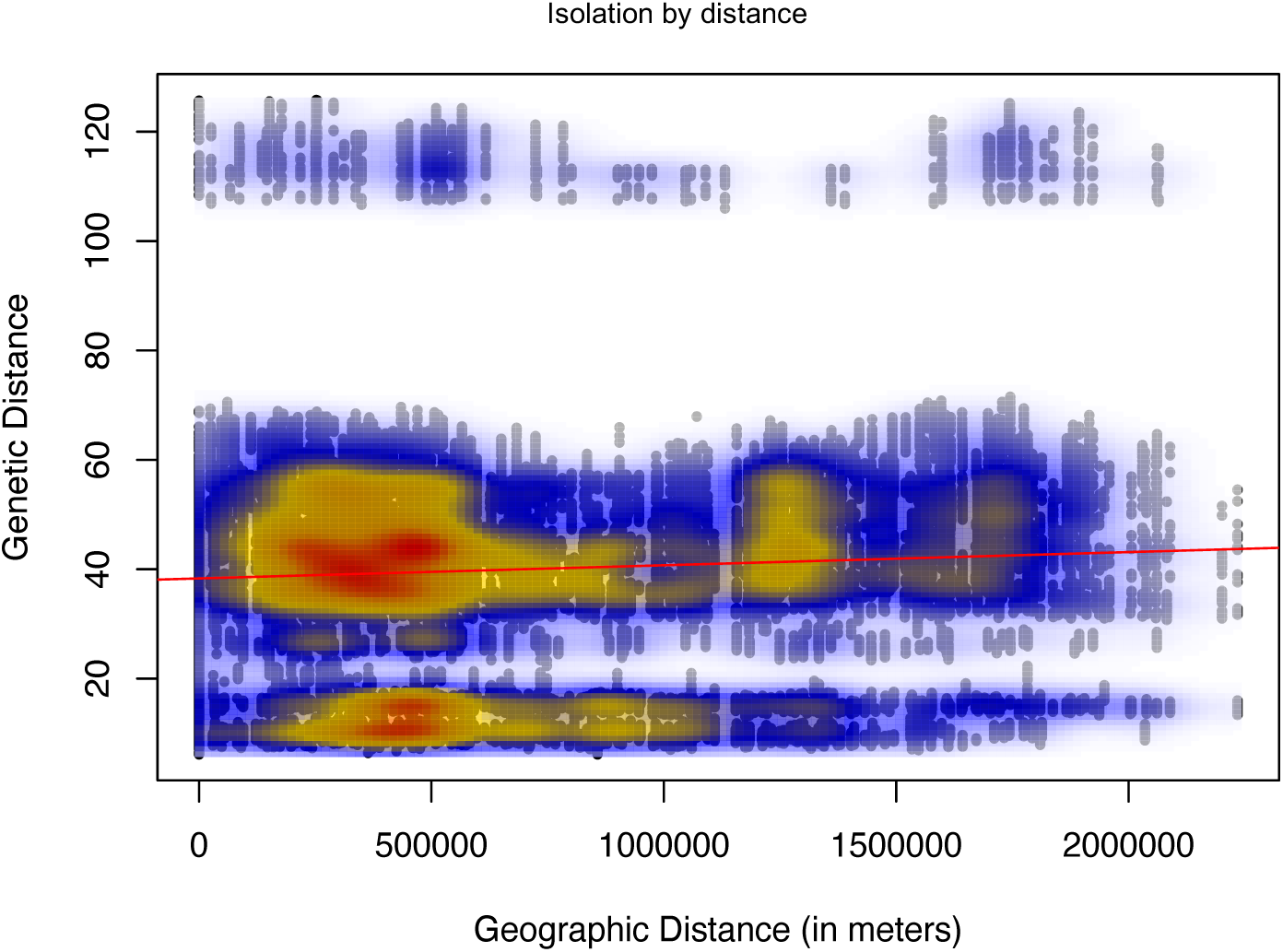
Isolation by distance analysis showing a weak correlation between genetic and geographic distance.

**Figure 5.**
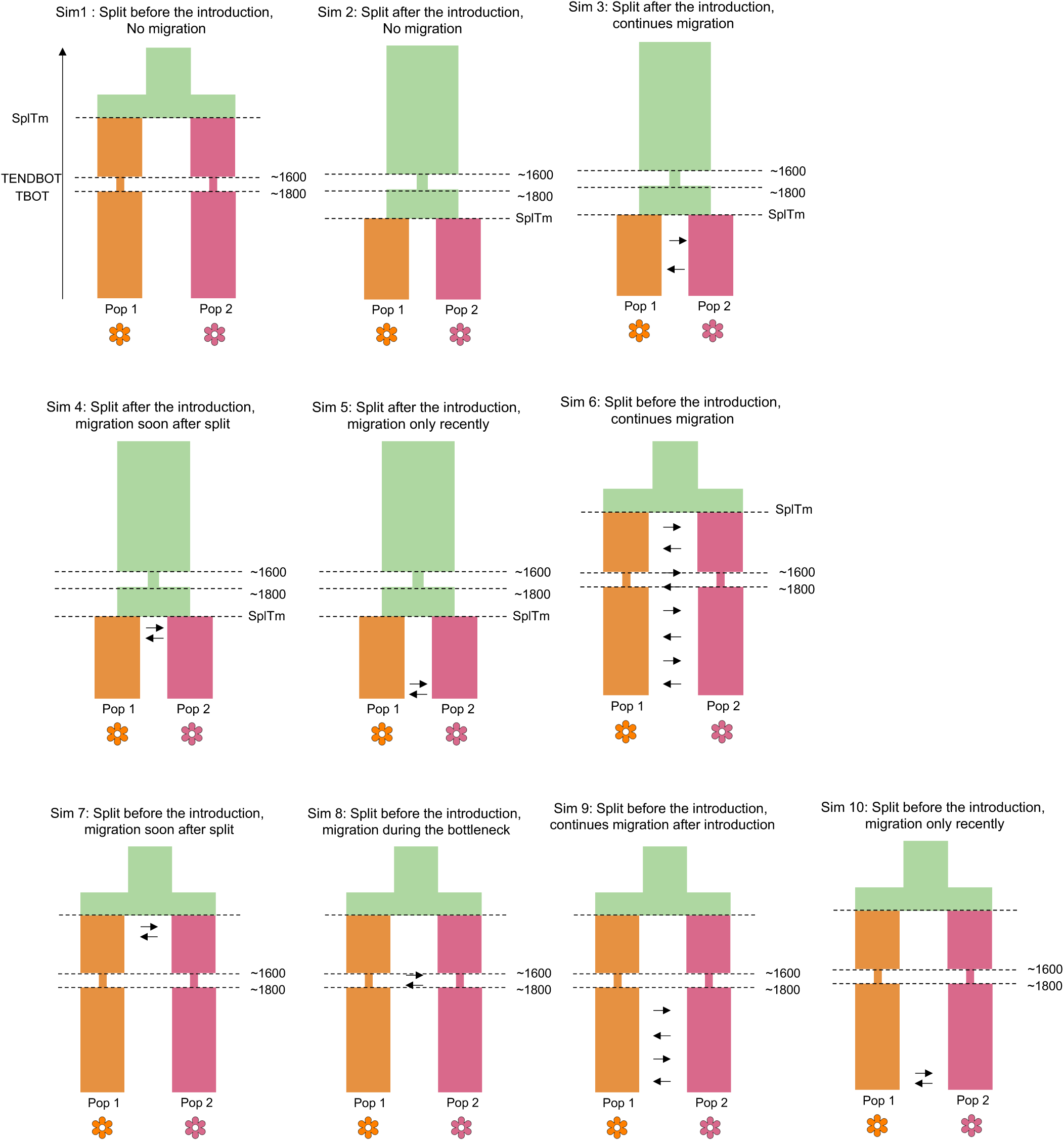
Fastsimcoal2 simulation scenarios. Different flower-colour lineages were simulated with divergence occurring either before or after the introduction, with or without migration. The period approximately from 1600 to 1800 represents the bottleneck associated with introduction to other regions. SplTm denotes the split time, while arrows indicate migration events. TBOT and TENDBOT represent the start and end of the bottleneck, respectively.

**Figure 6.**
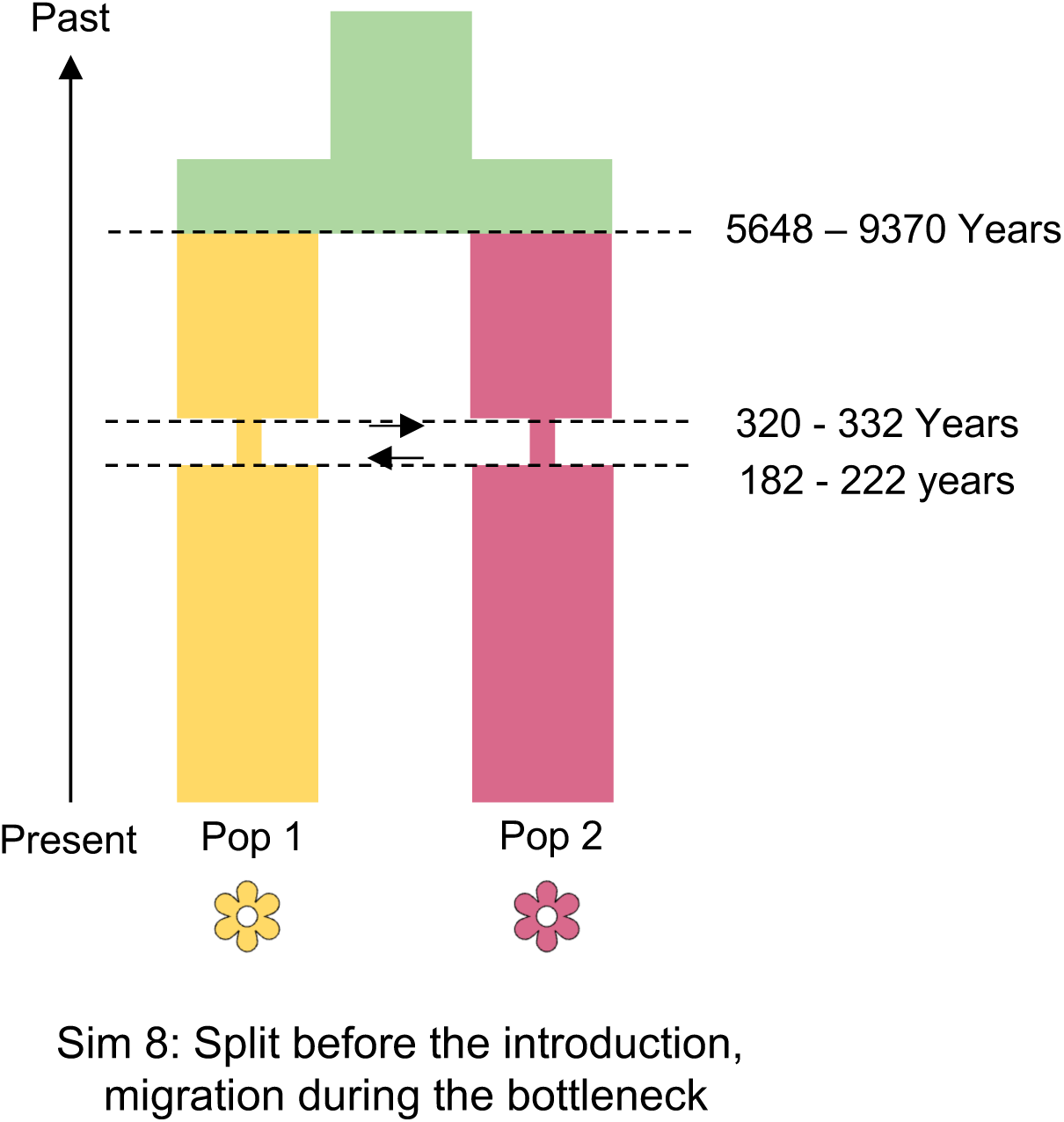
Selected Fastsimcoal scenario for yellow-pink and white-pink.

**Figure 7.**
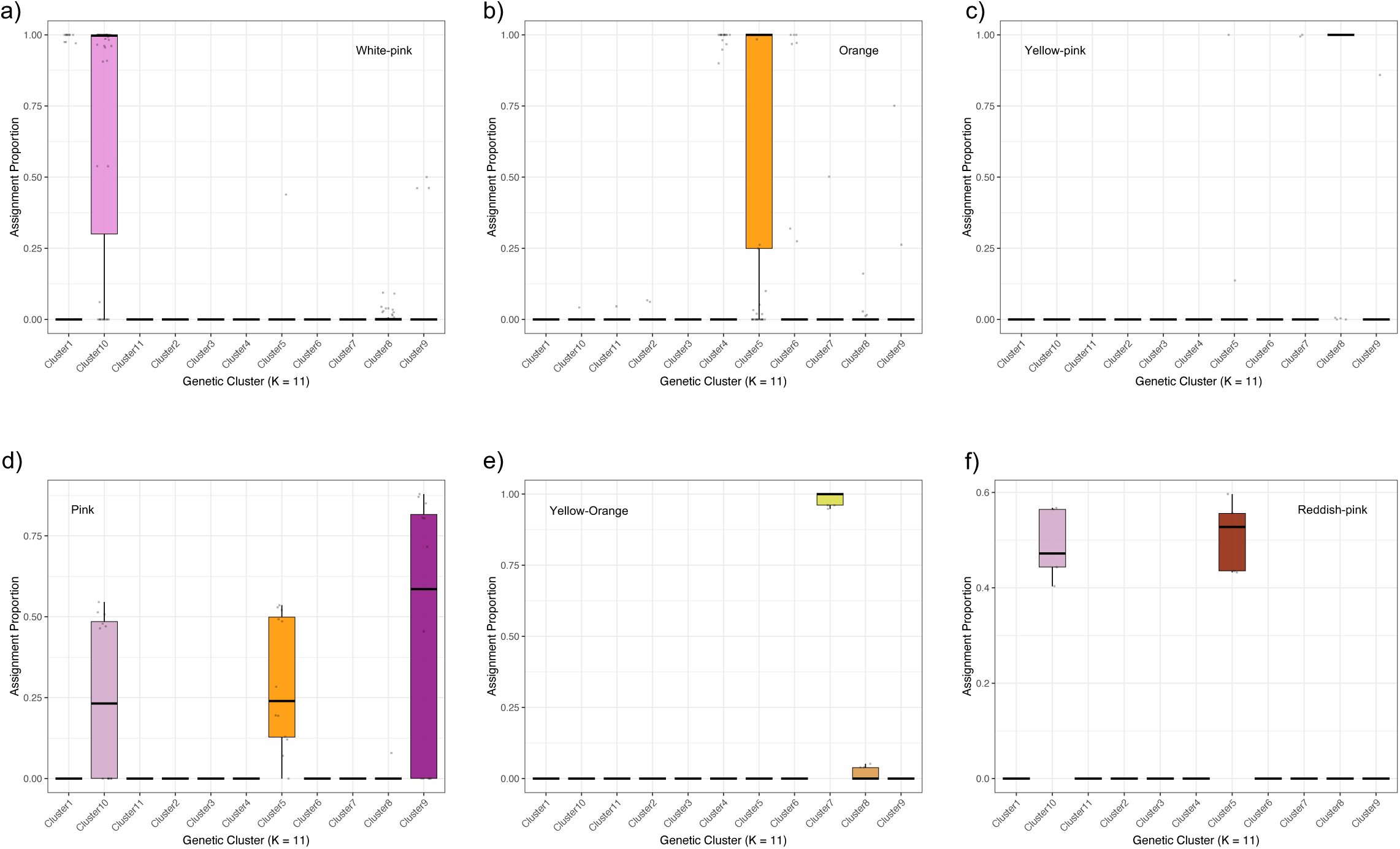
Proportions of cluster assignment for each flower colour type at K = 11. Cluster assignments are shown for the different flower colour types: (a) white–pink, (b) orange, (c) yellow–pink, (d) pink, (e) yellow–orange, and (f) reddish–orange. The x-axis represents clusters, and the y-axis represents assignment proportions.

**Table 1.**
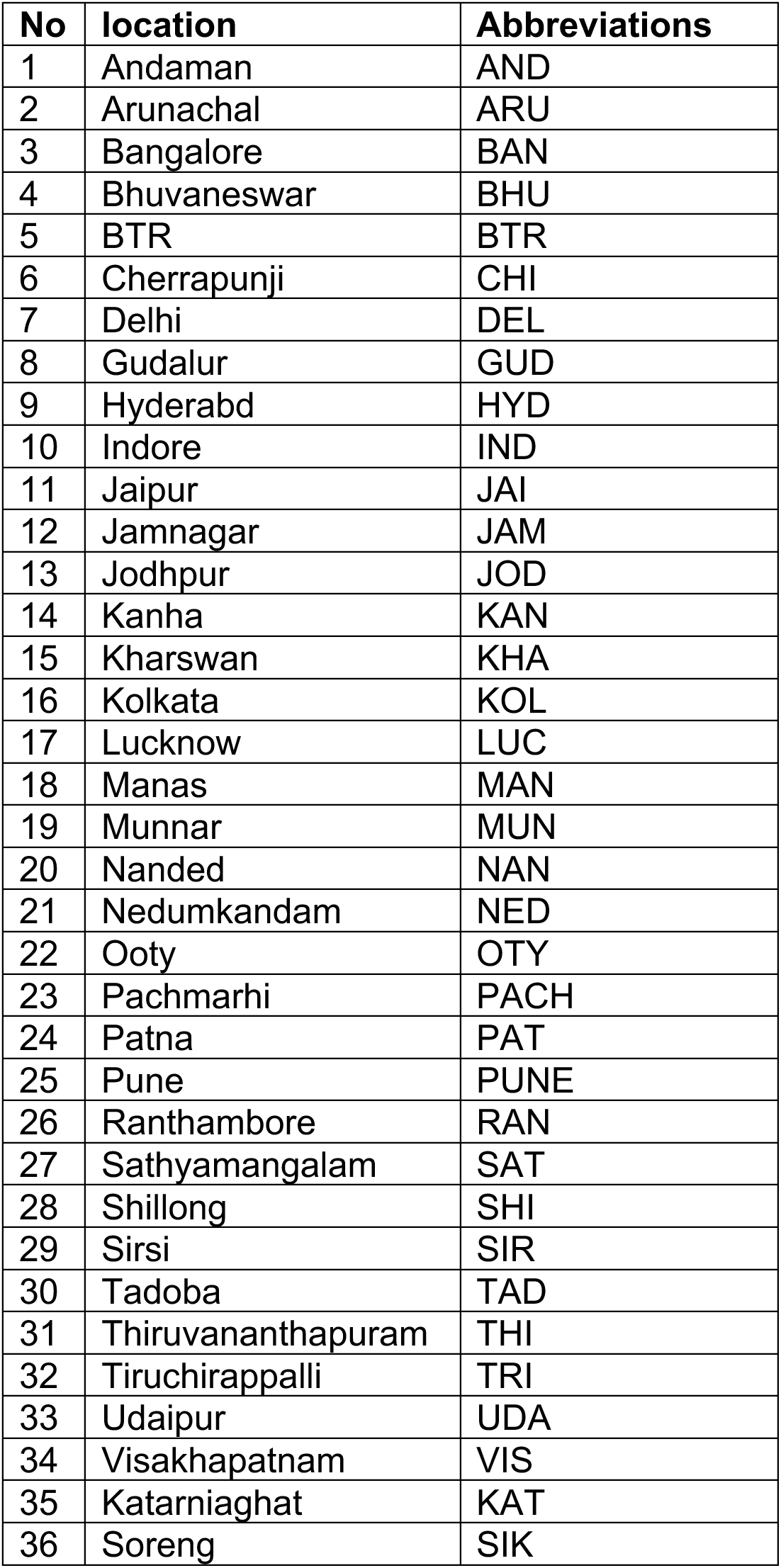
Sampling locations.

**Table 2.**
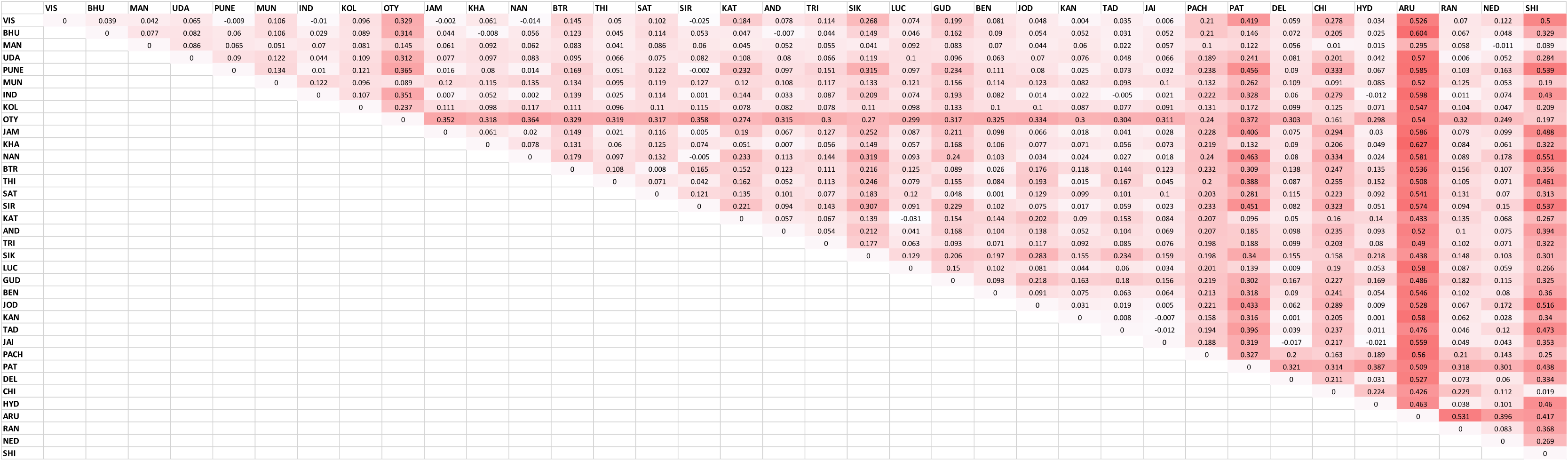
F_ST_ between different populations of Lantana in India.

**Table 3.**
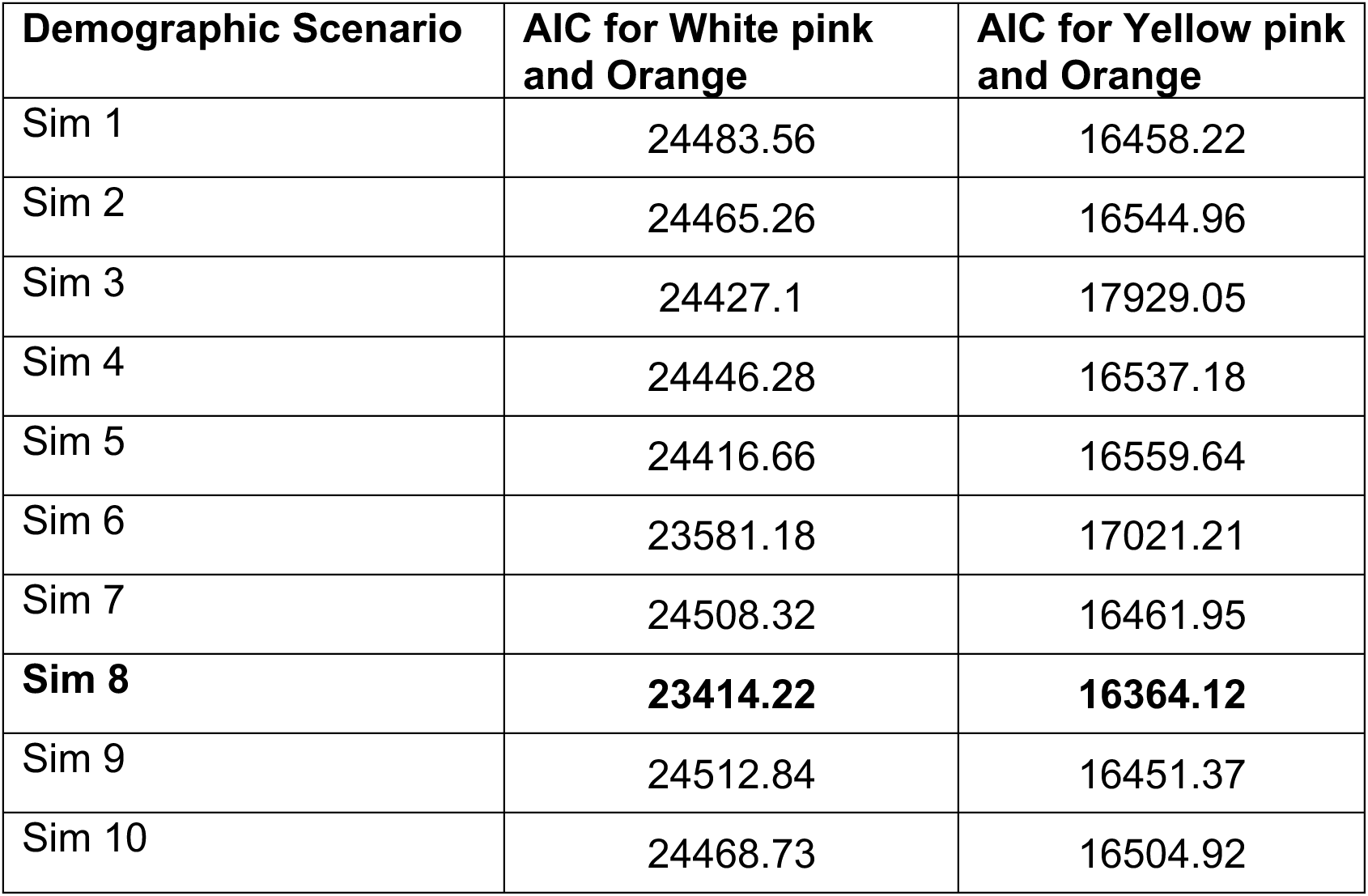
AIC values for alternative demographic scenarios tested for both flower colour combinations.

**Table 4.**
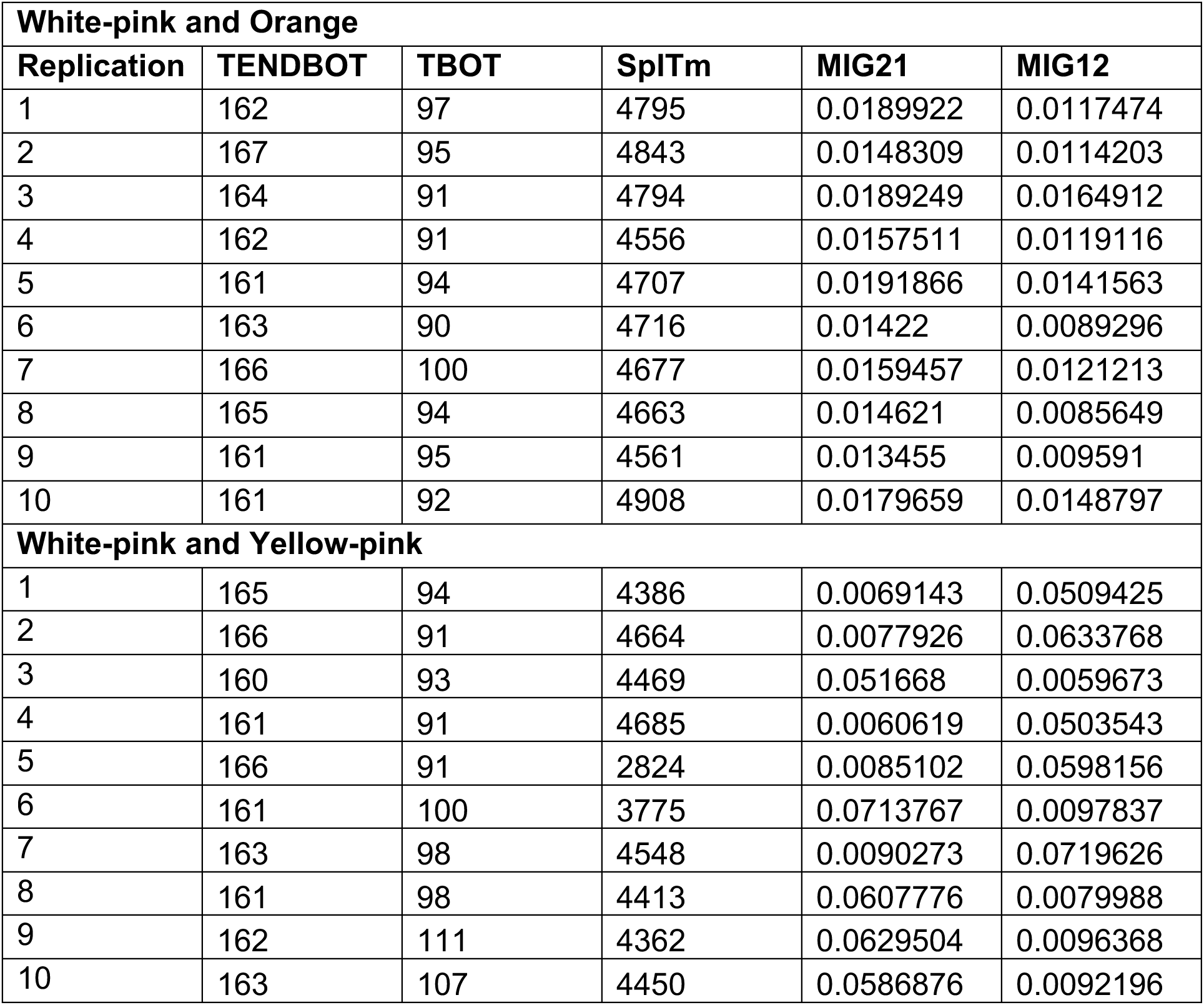
Estimated demographic parameters under the Sim8 scenario for the orange and white–pink morphs. TBOT – Start of bottleneck backward in time, TENBOT – end of bottleneck, SplTm – Split time, MIG21 – Migration from 2 to 1, MIG12 – Migration form 1 to 2.

**Table 5.**
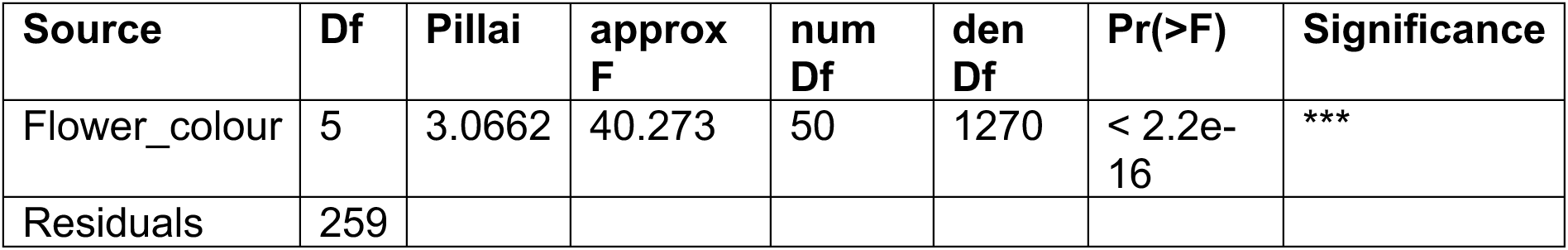
Summary statistics for the MANOVA.

