## Supplementary information for "The population structure of invasive *Lantana camara* is shaped by its mating system"

**Supplementary material of the paper “*The population structure of invasive Lantana camara is shaped by its mating system”***

P. Praveen, Rajesh Gopal, Uma Ramakrishnan

Table 1 Sampling locations

| **No** | **location** | **Abbreviations** |
| --- | --- | --- |
| 1 | Andaman | AND |
| 2 | Arunachal | ARU |
| 3 | Bangalore | BAN |
| 4 | Bhuvaneswar | BHU |
| 5 | BTR | BTR |
| 6 | Cherrapunji | CHI |
| 7 | Delhi | DEL |
| 8 | Gudalur | GUD |
| 9 | Hyderabd | HYD |
| 10 | Indore | IND |
| 11 | Jaipur | JAI |
| 12 | Jamnagar | JAM |
| 13 | Jodhpur | JOD |
| 14 | Kanha | KAN |
| 15 | Kharswan | KHA |
| 16 | Kolkata | KOL |
| 17 | Lucknow | LUC |
| 18 | Manas | MAN |
| 19 | Munnar | MUN |
| 20 | Nanded | NAN |
| 21 | Nedumkandam | NED |
| 22 | Ooty | OTY |
| 23 | Pachmarhi | PACH |
| 24 | Patna | PAT |
| 25 | Pune | PUNE |
| 26 | Ranthambore | RAN |
| 27 | Sathyamangalam | SAT |
| 28 | Shillong | SHI |
| 29 | Sirsi | SIR |
| 30 | Tadoba | TAD |
| 31 | Thiruvananthapuram | THI |
| 32 | Tiruchirappalli | TRI |
| 33 | Udaipur | UDA |
| 34 | Visakhapatnam | VIS |
| 35 | Katarniaghat | KAT |
| 36 | Soreng | SIK |

*Table 2 F_ST_ between different populations of Lantana in India*

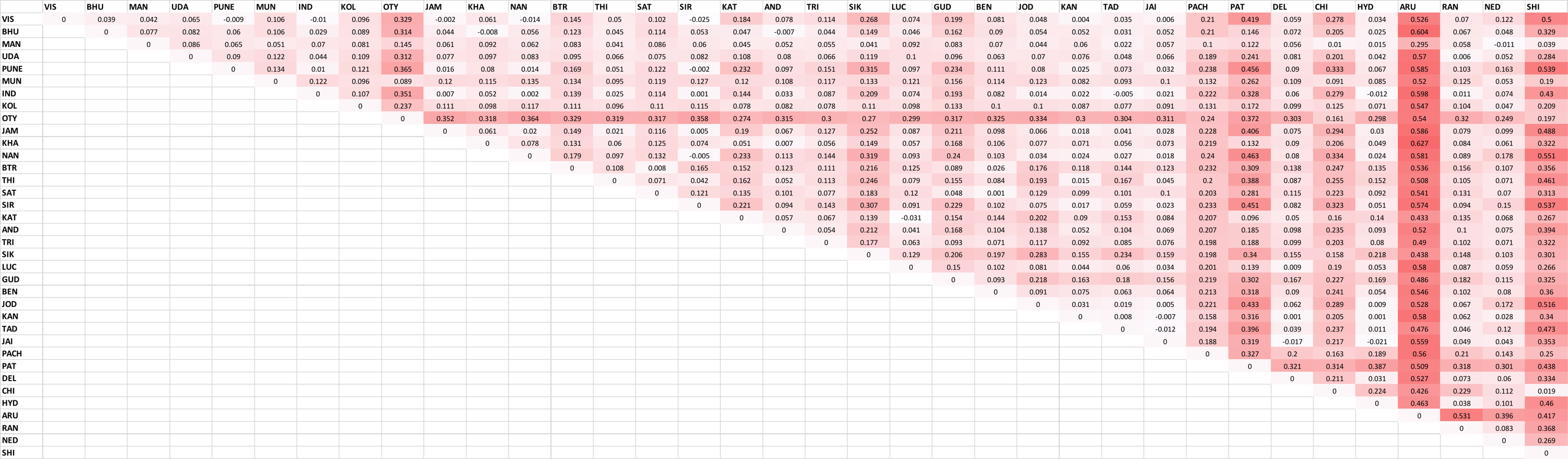

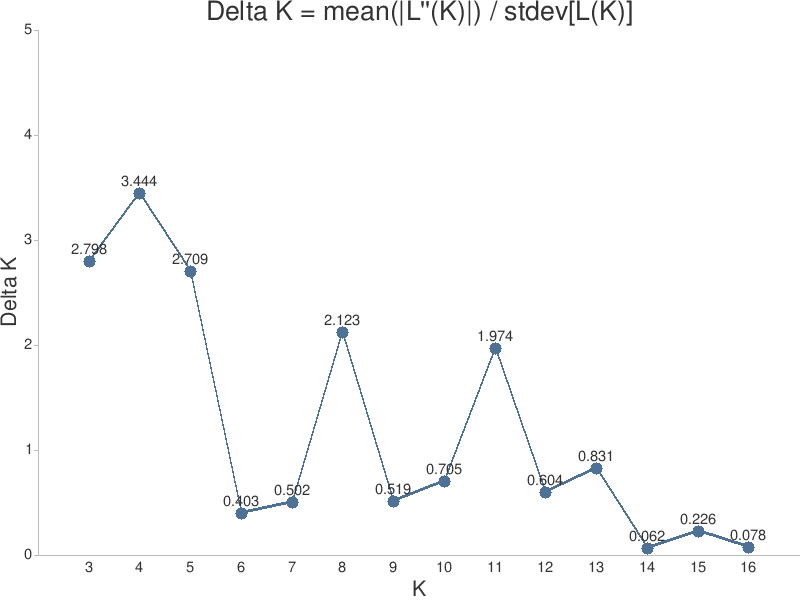

Delta K

K

Figure 1 Optimum number of clusters using Evanno’s delta K method

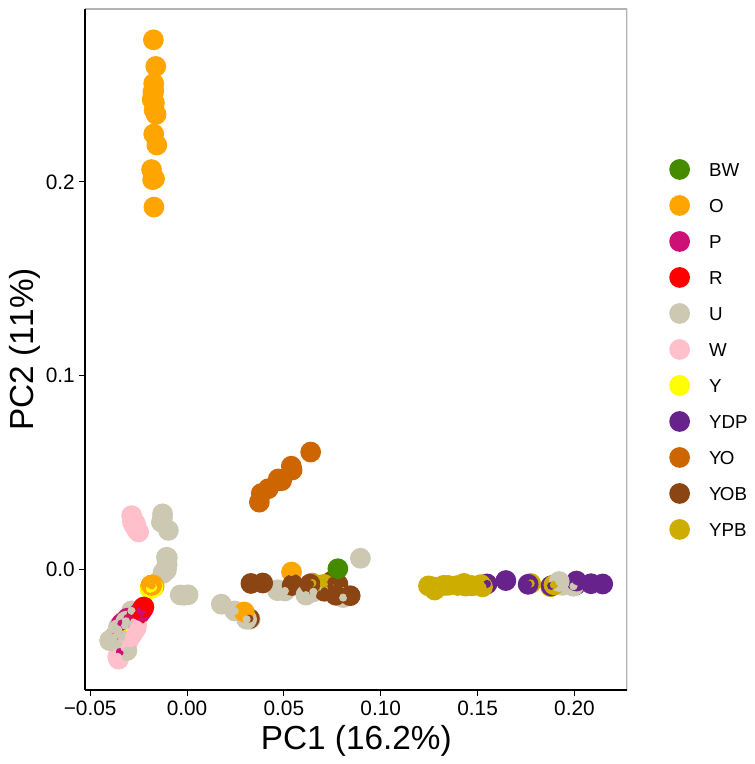

White lantana

Orange

Pink-pink

Reddish-pink

Unknown

White-pink

Yellow-pink -a

Yellow-dark pink

Yellow-orange – a

Yellow-orange – b

Yellow-pink - b

*Figure 2 PCA based on flower colour of Lantana individuals in India. Individuals with similar flower colours are coloured similarly in the plot. The flower colour of some of the samples was unknown.*

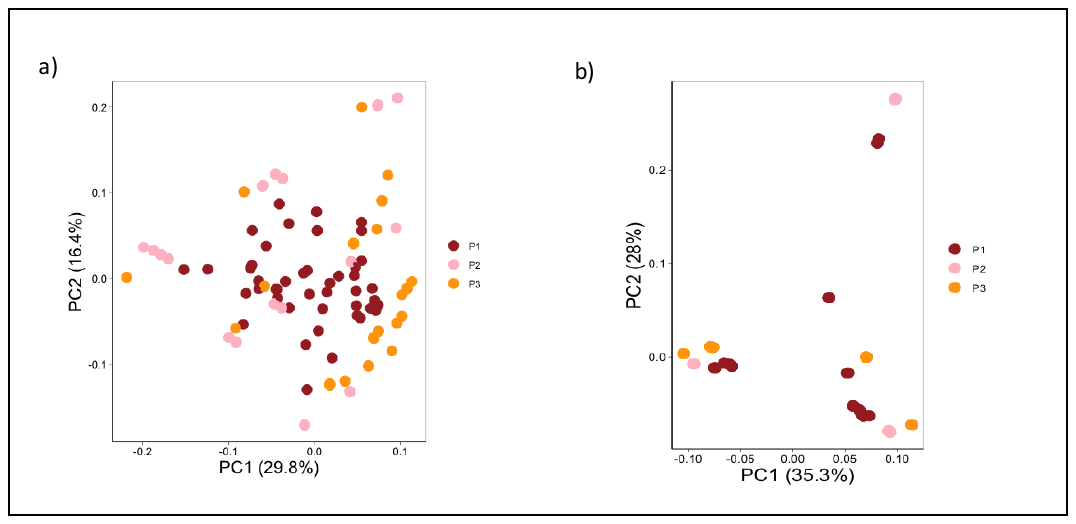

Figure 3 PCA of 50 randomly selected individuals from the simulation scenarios. P1 is the simulated native population and P2 and P3 are the simulated invasive populations. a) PCA of cross-fertilizing individuals simulated b) PCA of self-fertilizing individuals simulated

Isolation by distance

*Figure 4 Isolation by distance analysis showing a weak correlation between genetic and geographic distance*

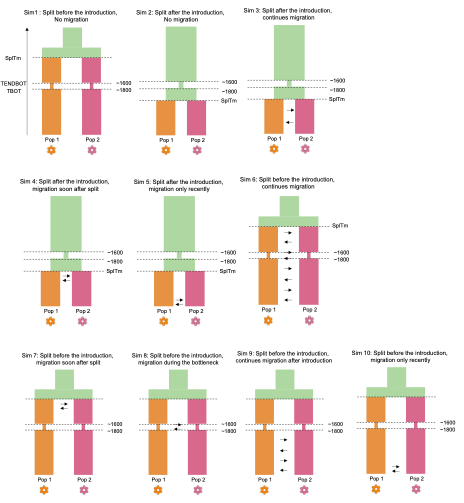

Figure 5 Fastsimcoal2 simulation scenarios. Different flower‐colour lineages were simulated with divergence occurring either before or after the introduction, with or without migration. The period approximately from 1600 to 1800 represents the bottleneck associated with introduction to other regions. SplTm denotes the split time, while arrows indicate migration events. TBOT and TENDBOT represent the start and end of the bottleneck, respectively.

Table 3 AIC values for alternative demographic scenarios tested for both flower colour combinations

| **Demographic Scenario** | **AIC for White pink and Orange** | **AIC for Yellow pink and Orange** |
| --- | --- | --- |
| Sim 1 | 24483.56 | 16458.22 |
| Sim 2 | 24465.26 | 16544.96 |
| Sim 3 | 24427.1 | 17929.05 |
| Sim 4 | 24446.28 | 16537.18 |
| Sim 5 | 24416.66 | 16559.64 |
| Sim 6 | 23581.18 | 17021.21 |
| Sim 7 | 24508.32 | 16461.95 |
| **Sim 8** | **23414.22** | **16364.12** |
| Sim 9 | 24512.84 | 16451.37 |
| Sim 10 | 24468.73 | 16504.92 |

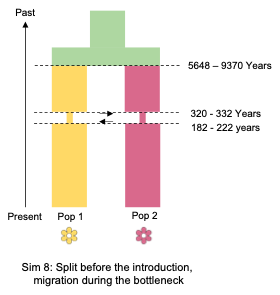

Figure 6 Selected Fastsimcoal scenario for yellow-pink and white-pink

Table 4 Estimated demographic parameters under the Sim8 scenario for the orange and white–pink morphs. TBOT – Start of bottleneck backward in time, TENBOT – end of bottleneck, SplTm – Split time, MIG21 – Migration from 2 to 1, MIG12 – Migration form 1 to 2

| **White-pink and Orange** | | | | | |
| --- | --- | --- | --- | --- | --- |
| **Replication** | **TENDBOT** | **TBOT** | **SplTm** | **MIG21** | **MIG12** |
| 1 | 162 | 97 | 4795 | 0.0189922 | 0.0117474 |
| 2 | 167 | 95 | 4843 | 0.0148309 | 0.0114203 |
| 3 | 164 | 91 | 4794 | 0.0189249 | 0.0164912 |
| 4 | 162 | 91 | 4556 | 0.0157511 | 0.0119116 |
| 5 | 161 | 94 | 4707 | 0.0191866 | 0.0141563 |
| 6 | 163 | 90 | 4716 | 0.01422 | 0.0089296 |
| 7 | 166 | 100 | 4677 | 0.0159457 | 0.0121213 |
| 8 | 165 | 94 | 4663 | 0.014621 | 0.0085649 |
| 9 | 161 | 95 | 4561 | 0.013455 | 0.009591 |
| 10 | 161 | 92 | 4908 | 0.0179659 | 0.0148797 |
| **White-pink and Yellow-pink** | | | | | |
| 1 | 165 | 94 | 4386 | 0.0069143 | 0.0509425 |
| 2 | 166 | 91 | 4664 | 0.0077926 | 0.0633768 |
| 3 | 160 | 93 | 4469 | 0.051668 | 0.0059673 |
| 4 | 161 | 91 | 4685 | 0.0060619 | 0.0503543 |
| 5 | 166 | 91 | 2824 | 0.0085102 | 0.0598156 |
| 6 | 161 | 100 | 3775 | 0.0713767 | 0.0097837 |
| 7 | 163 | 98 | 4548 | 0.0090273 | 0.0719626 |
| 8 | 161 | 98 | 4413 | 0.0607776 | 0.0079988 |
| 9 | 162 | 111 | 4362 | 0.0629504 | 0.0096368 |
| 10 | 163 | 107 | 4450 | 0.0586876 | 0.0092196 |

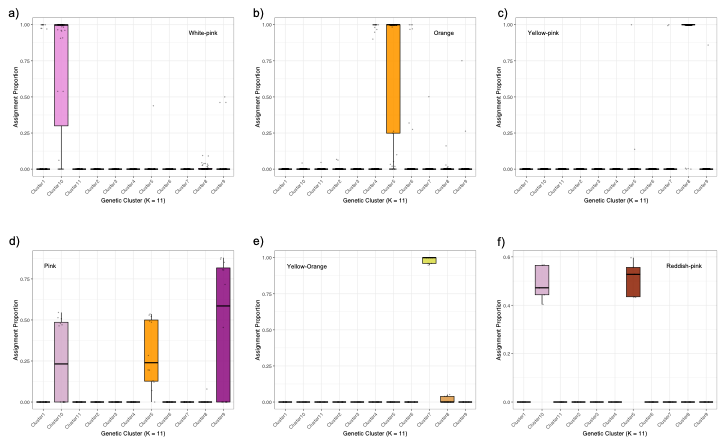

Figure 7 Proportions of cluster assignment for each flower colour type at K = 11. Cluster assignments are shown for the different flower colour types: (a) white–pink, (b) orange, (c) yellow–pink, (d) pink, (e) yellow–orange, and (f) reddish–orange. The x-axis represents clusters, and the y-axis represents assignment proportions.

Table 5 Summary statistics for the MANOVA

| **Source** | **Df** | **Pillai** | **approx F** | **num Df** | **den Df** | **Pr(>F)** | **Significance** |
| --- | --- | --- | --- | --- | --- | --- | --- |
| Flower_colour | 5 | 3.0662 | 40.273 | 50 | 1270 | < 2.2e-16 | *** |
| Residuals | 259 |  |  |  |  |  |  |
